# Liver single-nucleus multiome profiling reveals cell-type mechanisms for cardiometabolic traits

**DOI:** 10.1101/2025.08.06.668997

**Authors:** Abdalla A Alkhawaja, Kevin W Currin, Hannah J Perrin, Swarooparani Vadlamudi, Amy S Etheridge, K Alaine Broadaway, Gabrielle H Cannon, Carlton W Anderson, Anne H Moxley, Alina C Iuga, Erin G Schuetz, Federico Innocenti, Terrence S Furey, Karen L Mohlke

## Abstract

The liver is a central regulator of cardiometabolic physiology, coordinating processes such as lipid and glucose metabolism, protein synthesis, and detoxification. Genome-wide association studies (GWAS) have identified hundreds of genetic variants associated with cardiometabolic traits, yet their molecular mechanisms in liver cell types remain unclear. Using multiome single-nucleus RNA and ATAC sequencing on liver samples from 39 individuals, we profiled gene expression and chromatin accessibility in 68,398 nuclei across six primary liver cell types. We identified 306,706 accessible chromatin regions, including 70,884 regions undetected in bulk tissue analyses and that predominantly represent less abundant cell types. To identify genetic effects on gene regulation in liver cell types, we mapped quantitative trait loci (QTLs) and detected 1,885 chromatin accessibility QTLs (caQTLs) and 67 expression QTLs (eQTLs). We integrated cell-type QTLs with GWAS signals and revealed cell types, genes, and chromatin regulatory elements involved in cardiometabolic traits, such as liver enzyme and cholesterol levels. Non-hepatocyte cell-type QTL analyses exposed previously obscured mechanisms, such as an eQTL for *ADAMTS12* in liver sinusoidal endothelial cells potentially involved in liver fibrosis, demonstrating that single-nucleus approaches can capture regulatory events missed in bulk analyses. Furthermore, we predicted cell type of action for bulk liver caQTLs colocalized with GWAS signals, enhancing mechanistic insights for complex trait associations. Our findings provide a high-resolution map of the hepatic regulatory landscape and advance the understanding of cellular contexts and molecular mechanisms underlying cardiometabolic traits.

## Introduction

Genome-wide association studies (GWAS) have identified hundreds of signals for cardiometabolic traits^1–6^, but the molecular mechanisms and cell type(s) of action for most of these signals have not yet been defined. GWAS signals are enriched in regulatory elements of trait-relevant tissues^7–9^. Studies of quantitative trait loci for gene expression (eQTLs)^10^ and chromatin accessibility (caQTLs)^11–14^ have identified GWAS variants that may regulate gene expression and contribute to phenotypic variation. Profiles of gene expression and chromatin accessibility in single cells or nuclei have been used to predict specific functional variants and identify cell-type regulatory landscapes and target genes at many GWAS signals^15–19^. To annotate additional GWAS signals, single nucleus gene expression and chromatin accessibility profiles are needed in more disease-relevant tissues.

Liver tissue regulates many processes involved in cardiometabolic disease, including lipid and glucose metabolism, inflammation, protein synthesis, and drug detoxification^20,21^. Bulk liver tissue eQTLs^22,23^ and caQTLs^11,13^ have predicted functional variants, regulatory elements, and target genes at GWAS signals for liver-relevant traits. While differences in gene expression between liver cell types have been reported^24–27^, only one study reported cell type-level eQTLs in human liver^28^. In addition, fewer studies have analyzed liver cell-type chromatin accessibility^17,29^, and cell type-level caQTLs have not yet been mapped in human liver. Single nucleus gene expression and chromatin accessibility profiles from multiple individuals are needed to identify genetic effects on gene regulation in specific liver cell types.

Here, we jointly profiled chromatin accessibility and gene expression in 68,398 single nuclei from liver tissue of 39 individuals. We characterized the chromatin accessibility landscape of liver cell types and identified enrichment of GWAS signals in the cell-type chromatin profiles. Leveraging data across multiple individuals, we identified cell-type eQTLs and caQTLs and used them to predict mechanisms of GWAS signals. Finally, we used cell-type accessible chromatin regions to predict the cell types of action for GWAS-colocalized caQTLs from bulk liver tissue.

## Material and Methods

### Liver tissue

Human liver tissue was collected from 40 deceased organ donors through the National Institutes of Health Liver Tissue Cell Distribution System (LTCDS). Tissue was obtained from the LTCDS and approved for use in this study as non-human subjects research by the Institutional Review Boards (IRBs) at St Jude Children’s Research Hospital (Memphis, TN) and the University of North Carolina (Chapel Hill, NC). Fibrosis and steatosis were assessed using hematoxylin and eosin (H&E) and Masson’s trichrome-stained slides from frozen tissue samples fixed in neutral buffered formalin.

### Genotyping and imputation

Genotyping of DNA from the 40 donors in the current study on the Illumina Human610-Quad v1.0 BeadChip was previously described as part of a larger study of 224 liver donors^22^. We performed genotype imputation to the TOPMed reference panel version r2^30^ together with 117 additional liver donors as previously described^11^. We retained imputed genotypes with imputation r^2^ > 0.3 using BCFtools v1.15.1^31^. We inferred genetic similarity to major 1000 Genomes populations using genotype principal components (PCs) as previously described^11^.

### Nuclei isolation

We isolated nuclei from ∼12 mg of frozen human liver tissue for each sample. We pooled aliquots from sets of five individuals, pulverized using a cell crusher (CellCrusher), lysed cells in ice-cold lysis buffer (10 mM Tris-HCl [pH 7.4], 10 mM NaCl, 3 mM MgCl_2_, 1% BSA, 0.1% Tween-20, 0.1% Nonidet P40, 0.01% digitonin, 1mM dithiothreitol, 0.04 U/ul RNase inhibitor) with vigorous shaking for 5 min at 4°C, and homogenized using a 2-mL dounce with 10 strokes of the loose pestle followed by 30 strokes of the tight pestle. We centrifuged the homogenized samples at 500 x g for 5 min at 4°C, resuspended in 1 mL ice-cold wash buffer (10 mM Tris-HCl [pH 7.4], 10 mM NaCl, 3 mM MgCl_2_, 1% BSA, 0.1% Tween-20, 1 mM dithiothreitol, 0.04 U/ul RNase inhibitor), sequentially filtered through 70 um, 30 um, and 20 um filters, centrifuged at 500 x g for 5 min at 4°C, and resuspended in 300 ul wash buffer. We then washed nuclei suspensions twice by layering over 300 ul of 1.5 M sucrose and centrifuging at 1,000 x g for 10 min at 4°C. We resuspended final nuclei pellets in 1 mL wash buffer, centrifuged at 500 x g for 5 min at 4°C, and resuspended in 35 ul of diluted nuclei buffer (1X 10x Genomics Nuclei Buffer, 1 mM dithiothreitol, 1 U/ul RNase inhibitor).

### 10x multiome assays

We performed single-nucleus chromatin accessibility and gene expression profiling using the 10x Genomics Chromium Controller and Chromium Next GEM Single Cell Multiome ATAC + Gene Expression Reagent Bundle following the manufacturer’s instructions. Briefly, we stained nuclei with acridine orange and propidium iodide (Logos Biosystems PN-F23001) and assessed for viability, concentration, and singleness using a LUNA-FL Dual Fluorescence Cell Counter (Logos Biosystems). We loaded 20,000 nuclei per inlet on a Chromium Controller instrument (10x Genomics). We purified sequencing libraries using SPRIselect beads, visualized using the Bioanalyzer High Sensitivity DNA chip, and quantified via KAPA Library Quantification Kit for Illumina Platforms (Roche PN-KK4824). We separately pooled ATAC and RNA sequencing libraries, spiked with 1% PhiX sequencing control (Illumina), and sequenced on a NovaSeq 6000 S4 array at the UNC High Throughput Sequencing Facility in paired-end format to a total depth of ∼8 billion read pairs passing quality filters. We used CellRanger-ARC Count v2 (10X Genomics) to align sequencing reads to the GRCh38-2020-A human reference genome, using the filtered GENCODE V32^32^ GTF file for gene quantification.

### Identification of singlet droplets and donor deconvolution

To prepare genotype VCF files, we selected single nucleotide polymorphisms (SNPs) that overlapped gene coordinates from the Cell Ranger GTF file using BCFtools v1.15.1^31^ and BEDTools v2.29^33^. To prepare BAM files, we selected unique alignments (mapq=255 from STAR) using samtools v1.15.1^31^, selected alignments overlapping SNPs using BEDTools v2.29^33^, and reordered chromosomes to match the order in the VCF file using the ReorderSam tool from picard (http://broadinstitute.github.io/picard). We performed genotype-based donor deconvolution and singlet identification using demuxlet^34^ as implemented in popscle (https://github.com/statgen/popscle). We ran demuxlet separately per multiome well using barcodes with at least 1,000 RNA UMI and using parameters –field GT –min-BQ 20 –min-MQ 30 –min-mac 4 –min-callrate 0.95. To test for sample swaps between wells, we provided the full list of 40 donors to the –sm-list option for each demuxlet run. We retained barcodes that demuxlet classified as singlets and assigned them to the best matched donor.

### Nuclei filtering and cell type annotation

We applied several quality control (QC) filters to select high quality nuclei from the set of singlet barcodes. For RNA, we selected barcodes with ≥1,000 UMI, ≥500 expressed genes, and ≤10% of reads mapping to the mitochondrial chromosome using Seurat v5^35^. For ATAC, we counted reads into a set of bulk liver tissue ATAC peaks mapped in 138 individuals^11^ and selected barcodes with >500 fragments in peaks, >15% fragments overlapping peaks, and transcription start site (TSS) enrichment >2 (calculated from EnsDb.Hsapiens.v86) using Signac^36^.

We used DecontX^37^ to minimize ambient RNA contamination. To generate a set of cell-type labels, we clustered nuclei by their gene expression profiles separately per batch (set of nuclei pooled from 5 individuals) using Seurat^35^. We log-normalized gene counts, calculated PCs using the 2,000 most variable genes, constructed shared nearest neighbor graphs using the first 20 PCs, identified clusters using the Louvain algorithm (resolution = 0.25), and generated UMAP embeddings using the 20 PCs. We assigned clusters to cell types based on established cell-type markers^27^. We observed multiple hepatocyte clusters, which we collapsed to a single cell type. We then ran DecontX separately by batch using these cell types and defining barcodes with ≤100 UMI as background. Next, we re-applied the gene expression filters, which resulted in 39 samples with nuclei that passed QC.

Following decontamination, we merged corrected count matrices from all batches and identified clusters. To account for batch effects, we calculated 50 PCs from the decontaminated RNA data and used Harmony^38^ as implemented in the Seurat IntegrateLayers function. Next, we recalculated clusters and UMAP projections, and annotated clusters. To obtain more granular cell type annotation, we used Azimuth (https://azimuth.hubmapconsortium.org/) in R with the supplied liver dataset as the reference to transfer annotations to our gene expression data.

### Cell-type accessible chromatin regions

We used the barcodes of the gene expression-based annotated nuclei to subset the Cell Ranger ATAC-seq BAM files by cell type and donor combination and generate pseudobulk BAM files. We selected non-duplicate reads with high mapping quality to primary nuclear chromosomes using samtools v1.17^31^ with parameters -f 3 -F 4 -F 8 -F 256 -F 2048 -F 1024 -q 30. Due to low read depth, we merged BAM files across donors within a cell type using samtools^31^ prior to peak calling. We then converted the merged BAM files to BED using BEDTools v2.29^33^ and called peaks for each cell type using MACS2^39^ v2016-02-15 with parameters –nomodel –shift -100 –extsize 200 -q 0.05 –keep-dup all; the –keep-dup all option was used because duplicate removal at the nucleus level was previously performed using samtools and we wanted to retain reads mapping to the same genomic location but in different nuclei. We removed peaks overlapping ENCODE exclusion regions^40^ using BEDTools^33^. We used ataqv v1.0.0^41^ to calculate quality metrics of the pseudobulk datasets, including transcription start site (TSS) enrichment and percentage of reads overlapping peaks. We generated a single peak set by merging peaks across cell types using BEDTools^33^.

To cluster nuclei using their chromatin accessibility profiles, we normalized counts using the TF-IDF function and used all peaks to calculate 50 latent semantic indexing (LSI) components. We computed shared nearest neighbor graphs, clusters, and UMAP projections as described for the gene expression analysis, excluding the first LSI component as it correlated with library size. To account for batch effects, we ran Harmony^38^ on 50 LSI components. To integrate the gene expression and chromatin accessibility UMAP projections, we computed weighted nearest neighbors (WNN) using 50 gene expression PCs and 49 chromatin accessibility LSI components (excluding the first). We used the gene expression annotations for cell type identification.

To identify cell-type peaks that were missed in analyses of bulk tissue, we determined which cell-type peaks shared at least 1 basepair (bp) with previously reported bulk liver tissue ATAC-seq peaks^11^ using the plyranges R package^42^.

### Cell-type marker ATAC-seq peaks

We used differential chromatin accessibility analysis to identify marker peaks for each cell type. To compare hepatocyte nuclei to other cell types, we downsampled the number of hepatocyte nuclei in each donor to 10% using the sample function in R^43^ (random number generator seed of 123). We then called peaks on the downsampled hepatocytes and merged peaks across cell types using downsampled hepatocytes instead of full-depth hepatocytes. We counted the number of ATAC-seq reads from the pseudobulk BAM files that overlapped the merged cell-type peaks using featureCounts^44^, corrected for peak-level GC content using EDASeq^45^, and corrected for library size using DESeq2^46^ size factors. We only tested peaks with a minimum mean count of 10 across donors in a cell type for marker peak status in that cell type. We tested for differentially accessible peaks between each pair of cell types using DESeq2^46^ and defined marker peaks as peaks more accessible (FDR<5%, Benjamini-Hochberg (BH) procedure^47^) in one cell type compared to all other cell types.

We used the Genomic Regions Enrichment of Annotations Tool (GREAT)^48^ to test if marker peaks were located near genes involved in cell type-relevant biological functions using the GO Biological Process ontology^49^ and using all merged cell type peaks as background. We tested if marker peaks were enriched for known transcription factor (TF) motifs using HOMER v4.11^50^ with the -size 200 parameter and using all merged cell type peaks as background.

### Heritability enrichment in cell-type peaks

We downloaded GWAS summary statistics from the Benjamin Neale lab analysis of the UK Biobank dataset (https://nealelab.github.io/UKBB_ldsc/downloads.html). We tested for heritability enrichment for traits with heritability z score ≥ 4 in cell type ATAC peaks using stratified LD score regression implemented in the LDSC software^9,51^. We ran a single LDSC model that consisted of 7 annotations: annotations for peaks in each of our identified 6 cell types (excluding B cells) and the baseline v1.2 annotation provided with LDSC. To run LDSC, we used liftOver^52^ to convert cell type peak coordinates to GRCh37, used the GRCh37-based LDSC reference files derived from 1000G phase 3 European genotypes, and performed analysis using HapMap SNPs. For each combination of trait and cell-type peak annotation, we calculated the p-value of the LDSC coefficient z-score using a one-sided test assuming a standard normal distribution. We calculated FDR of the coefficient p-values using the BH procedure^47^ and classified results with FDR<5% as significant.

### Mapping caQTL

We removed ATAC-seq reads exhibiting allelic mapping bias from the pseudobulk BAM files per cell type and donor using the remapping pipeline in WASP v0.3.4^53^ with updates as of February 9, 2023. Because the Cell Ranger alignment pipeline cannot be run in the WASP pipeline, we approximated the Cell Ranger alignment command using BWA-mem v0.7.17^54^ with parameters -M -I 250,150. For each cell type, we generated a peak-by-donor count matrix using featureCounts^44^, corrected for peak-level GC content using EDASeq^45^, and corrected for library size and performed variance stabilization using DESeq2^46^. We performed PCA on variance-stabilized peak counts and found that PC values were strongly driven by the fraction of 0 counts in each sample. Therefore, we did not use ATAC-seq PCs as covariates in the caQTL model. Instead, we used the first two genotype PCs, sex, and ATAC TSS enrichment as caQTL covariates. We only tested peaks for caQTL if the peaks had at least 5 counts in at least 10 donors for a given cell type. We used FastQTL^55^ to test for association between inverse normal-transformed peak counts and biallelic variants within 1 kilobase (kb) of peak centers and with minor allele frequency (MAF) of at least 0.1 in the 39 donors that passed QC. We corrected the FastQTL beta-adjusted p-values for genome-wide multiple testing using the BH procedure and defined peaks with a significant caQTL (FDR<5%) as caPeaks.

We classified a cell-type caPeak as shared with a caPeak in bulk liver tissue^11^ if the peaks overlapped by at least 1 bp and if their lead variants were in linkage disequilibrium (LD r^2^≥0.5, TOPMed^30^ Europeans calculated by TOPLD^56^).

### Mapping eQTL

To prepare the data for eQTL analysis, we generated pseudobulk expression profiles by summing the count matrices across cells for each cell type-donor combination using the AggregateExpression function from Seurat v5^35^. Per cell type, we kept genes with at least 10 counts in at least 10 donors. We corrected for library size and performed variance stabilization using DESeq2^46^. We used FastQTL^55^ to test for association of inverse normal-transformed gene counts and biallelic variants within 1 megabase (Mb) of the TSS of a gene and with MAF≥0.1. We included sex, two genotype PCs, and a variable number of expression PCs as covariates. Specifically, we included the first 2,3,6,2,3,6 PCs for hepatocytes, liver sinusoidal endothelial cells (LSECs), Kupffer cells , mesenchymal cells, NK-T cells and cholangiocytes respectively, which maximized the number of genes with significant eQTLs.

We adjusted the FastQTL beta-adjusted p-values for genome-wide multiple testing using the BH procedure and classified genes with a significant eQTL (FDR<5%) as an eGene. When multiple eQTLs were identified for the same eGene across cell types, we clumped eQTLs based on LD (r^2^ ≥ 0.2), which grouped 79 unclumped eQTLs into 67 distinct eQTLs.

We considered a cell-type eQTL to be shared with a bulk liver tissue eQTL from GTEx v8^10^ if the eGenes shared the same Ensembl ID or gene symbol and the lead variants were in strong LD (r^2^>0.5, TOPMed Europeans^30,56^). For cases in which the GTEx lead variant was not present in the TOPMed panel, we calculated LD between our cell-type lead variant and the GTEx second-best variant ranked by p-value.

### Identifying cell-type QTL shared with GWAS signals

We retrieved published GWAS summary statistics and signal lead variants for 17 cardiometabolic traits including lipids^1^, coronary artery disease^2^, plasma liver enzyme levels^3^, glycemic traits^4^, type 2 diabetes^5^, waist-hip ratio adjusted for body mass index^6^, BMI^57^, vitamin D^58^, and C-reactive protein^59^. We used summary statistics from individuals genetically similar to the European population for all GWAS studies. When studies did not provide conditionally distinct signals, we used those previously identified in our prior analyses^23^. To more comprehensively capture the LD structure of the secondary signal at *XKR9*, which lies at the edge of the locus boundary, we extended the signal window to 1 Mb upstream of the second signal lead variant before running the conditional analysis.

### Cell-type QTL shared with signals in the GWAS catalog

We retrieved signal lead variants from the NHGRI-EBI GWAS catalog^60^ (accessed June 4, 2025). For each trait, we retained variants with available rsID or genomic coordinates and selected those with the lowest p-value per trait. Cell-type eQTLs and caQTLs were considered to be shared with GWAS signals when LD r^2^ ≥ 0.5.

### Annotating bulk caQTL with cell-type peaks

We retrieved published bulk liver caQTL peaks and their predicted target genes and GWAS colocalizations^11^. The predicted target genes were linked to bulk liver caQTL peaks using four approaches: TSS proximity, caQTL – eQTL colocalization, promoter capture Hi-C, and shared caQTL signals for distal and promoter bulk peaks. We determined which cell-type peaks overlapped by at least 1 bp with bulk liver caQTL peaks using BEDTools intersect^33^. When plotting the example at the *GAS6* locus, we observed that the bulk liver ATAC peak (bulk peak94548) had two distinct bulk caQTL signals, thus we performed caQTL mapping for this peak including genotype dosage of one of them, rs74118417, as an additional covariate to the covariates used in the bulk study^11^ and used this conditioned signal in plotting.

## Results

### The cell-type regulatory landscapes of the human liver

To profile the regulatory elements and gene expression in cell types of the human liver, we performed single-nucleus ATAC (snATAC-seq) and RNA sequencing (snRNA-seq) on 40 genotyped liver samples (**Figure 1, Table S1**). After rigorous quality control, we identified 68,398 nuclei from 39 samples, averaging 1,753 nuclei per sample (**Figures S1, S2**). Using established cell-type gene expression markers^27^ (**Figure S3**), we identified seven broad cell types: hepatocytes, liver sinusoidal endothelial cells (LSECs), mesenchymal cells, Kupffer cells, NK-T cells, cholangiocytes, and B cells (**Figure 1**). Prior to batch correction, hepatocytes showed sample-specific clustering (**Figure S4-6**), consistent with earlier findings^24,27^. Although reference liver single-cell RNA-seq data^24–26,61,62^ enabled finer subtype classifications for non-parenchymal cells (**Figure S7**), we maintained broad cell-type definitions for downstream QTL analyses to maximize detection power. Cell-type proportions varied considerably between samples (**Figure 1**). Hepatocytes comprised the majority of nuclei, 68% (range: 36% - 85%, **Table S1**), while B cells were extremely sparse (1.5% of nuclei; **Figure 1**). Due to their low abundance, we excluded B cells from further analyses. These well-defined cell-type profiles lay the groundwork for understanding the cellular context of gene regulation in the liver.

**Figure 1:**
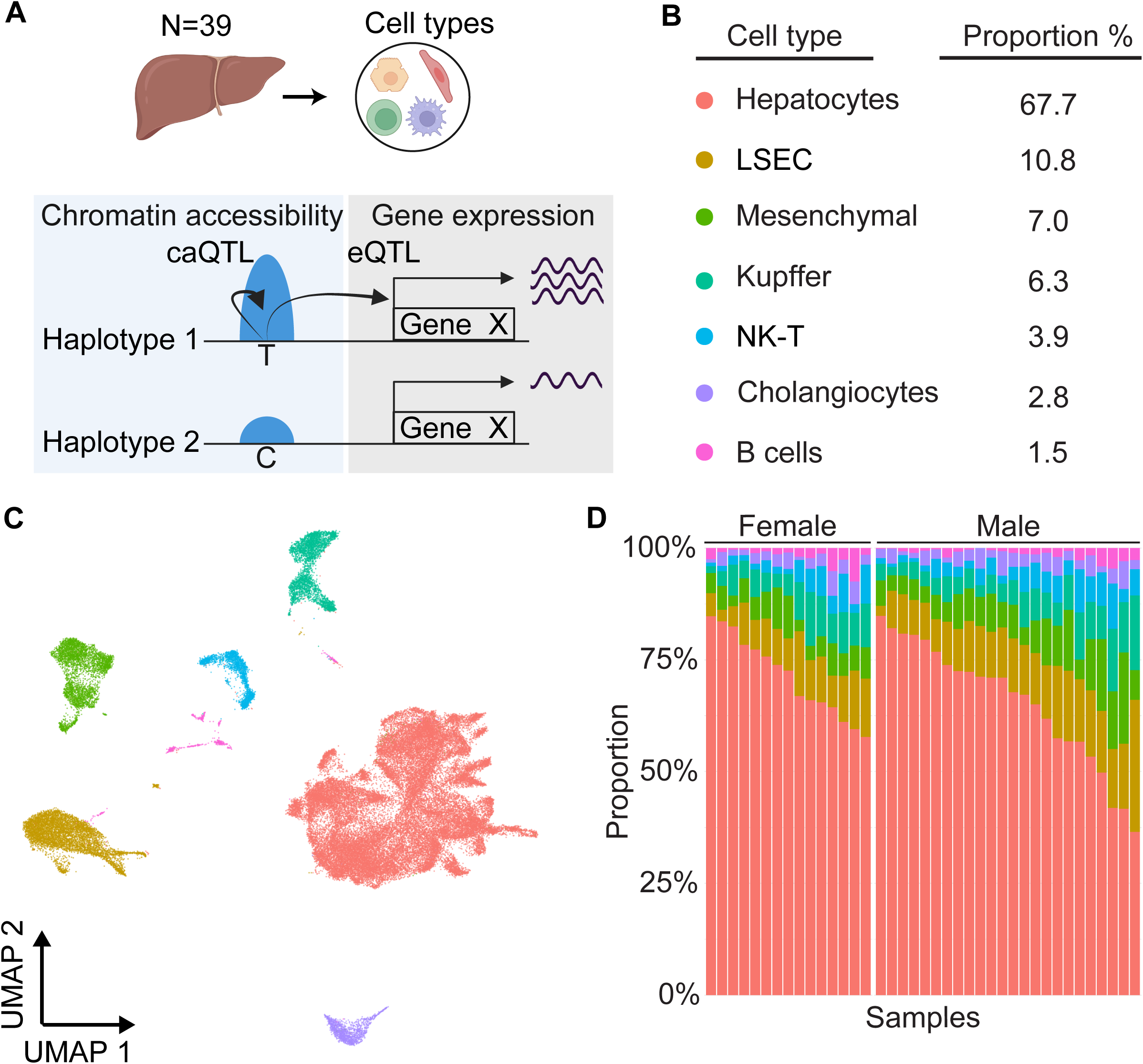
Identification and characterization of major cell types in human liver tissue using single-nucleus RNA & ATAC sequencing. (**A**) Project overview. Nuclei extracted from 39 human livers were sequenced to profile the chromatin and gene expression landscapes, identify eQTLs and caQTLs, and determine which signals are shared with GWAS. (**B**) Cell-type proportions of seven cell populations identified from snRNA-seq analysis of 68,393 nuclei. LSEC, liver sinusoidal endothelial cells. NKT, Natural killer T cells. (**C**) Joint UMAP of integrated snRNA-seq and snATAC-seq profiles showing seven main cell populations. Colors correspond to cell types shown in (B). (**D**) The cell-type composition of each sample, separated by sex. Colors correspond to cell types shown in (B).

Because each nucleus was profiled using snRNA-seq and snATAC-seq, the gene expression-based cell-type annotations apply to both modalities. To map the cell-type regulatory landscapes of the human liver, we identified 306,706 accessible chromatin regions (peaks) across the six cell types (**Supplemental data 1**). Cell clustering based on peak accessibility recapitulated gene expression-based cell-type clusters, indicating concordance between the two profiling methods (**Figure S5**). We identified 17,147 cell-type marker peaks that were more accessible in one cell type compared to all others (FDR < 0.05, **Figure S8**, **Table S2**, **Supplemental data 2**). These marker peaks were enriched near genes involved in canonical cell-type processes: lipid, carbohydrate, and drug metabolism genes for hepatocytes; angiogenesis and blood vessel morphogenesis genes for LSECs; and extracellular matrix organization genes for mesenchymal cells (**Figure S8**, **Table S3**). Consistent with these functional associations, marker peaks were enriched for binding motifs of known cell-type transcription factors (TFs), including HNF4A and the HNF1 family for hepatocytes, SPI1 for Kupffer cells, and SP1 for cholangiocytes (**Figure S8**, **Table S4**). Together these results demonstrate that we generated a high-quality cell type-resolved map of accessible chromatin profiles in human liver tissue.

To investigate the differences in regulatory element discovery between single nucleus and bulk ATAC-seq, we compared cell-type peaks to 349,685 bulk liver ATAC peaks from a larger cohort of 138 deeply sequenced individuals^11^. We found 235,822 cell-type peaks overlapped with bulk, while 70,884 peaks did not (**Figure 2**). As hepatocytes are the predominant cell type in the liver, we expected that most bulk peaks would originate from that cell type, and indeed 77% of cell-type peaks that were found in the bulk data were also detected in hepatocytes. In contrast, of the 70,884 cell-type peaks not found in the bulk analysis, only 27% were detected in hepatocytes (**Figure 2**, **supplemental data 1**), showing that cell-type ATAC-seq can identify peaks in less abundant cell types that are poorly represented in bulk ATAC-seq data. For example, a cell-type marker peak located in an intron of the Kupffer cell marker gene *LYN* was not detected in the bulk data nor hepatocytes (**Figure 2**). *LYN* is a member of the Src family of protein tyrosine kinases and plays a role in macrophage inflammatory response^63^. The 51,717 non-hepatocyte peaks not identified in the bulk analysis demonstrate the added resolution of snATAC-seq to uncover regulatory regions in less abundant liver cell types, enabling a more complete annotation of the hepatic regulatory landscape.

**Figure 2:**
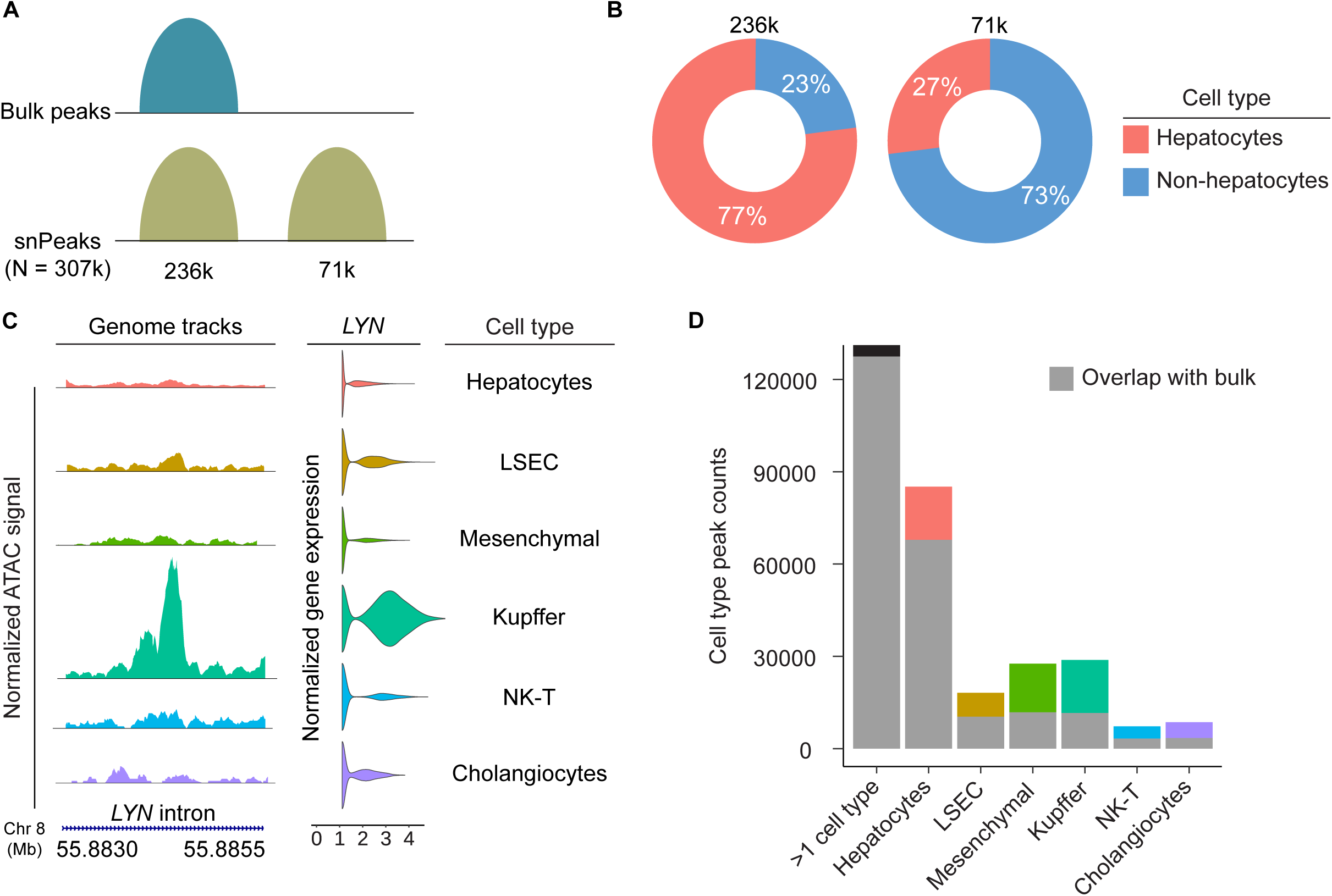
Accessible chromatin in liver cell types. (**A**) Among 307k accessible chromatin peaks observed in liver cell types using snATAC-seq, 236k overlapped accessible chromatin peaks detected in bulk liver ATAC-seq. (**B**) Among cell-type peaks that were detected in bulk liver ATAC-seq, only 23% were not detected in hepatocytes. Among peaks not detected in bulk liver ATAC-seq, 73% were not detected in hepatocytes. (**C**) An example of a cell-type marker peak in an intron of *LYN*. Coverage plot showing normalized ATAC-seq signal across cell types, with a Kupffer cell peak that did not pass thresholds of detection in other cell types or bulk liver tissue. The adjacent violin plots represent *LYN* gene expression by cell type. (**D**) Distribution of peaks detected in each cell type. Portions of bars shaded in gray represent peaks that were also detected in bulk liver. Portions of bars shaded in colors represent peaks only detected using snATAC-seq.

### Cell-type chromatin accessibility and gene expression QTLs

To investigate regulatory genetic variation across liver cell types, we conducted QTL analyses for both chromatin accessibility and gene expression. We identified variants significantly associated with chromatin accessibility for 1,885 peaks (caPeaks, FDR < 0.05), nearly entirely in hepatocytes (**Table 1**, **Table S5**). To confirm the robustness of these cell-type caQTL, we compared cell-type caPeaks with caPeaks detected in bulk liver tissue^11^: 90% overlapped with bulk caPeaks (**Table S6**). Cell-type and bulk caQTLs showed highly correlated effect sizes (Pearson’s R^2^ = 0.94, **Figure S9**). In parallel, we identified 67 eQTLs (FDR < 0.05) across the six liver cell types (**Table 1**, **Table S7**). We found 47 cell-type eGenes in common with bulk liver eQTLs in GTEx^10^, 38 of which (80%) shared the same eQTL signal (lead variants r^2^ ≥ 0.5, **Figure S9**, **Table S8**), such as the well-validated eQTL for *SORT1* detected in hepatocytes^64^. Together, these results underscore the reproducibility of the cell-type QTL data.

**Table 1:** Chromatin accessibility QTL (caQTL) and expression QTL (eQTL) identified in each cell type.

| **Cell type** | **caQTL count** | **eQTL count** |
| --- | --- | --- |
| Hepatocytes | 1880 | 55 |
| Endothelial | 4 | 8 |
| Kupffer | 1 | 7 |
| Mesenchymal | 0 | 4 |
| Cholangiocytes | 0 | 3 |
| NK-T | 0 | 2 |
| **Total** | **1885** | **79** |

Some eQTLs are called in >1 cell type. Clumping by LD $r^2 > 0.2$ results in 67 eQTLs.

Cell-type analysis offers the potential to identify eQTL signals in less abundant cell types. We investigated whether there were cell-type eQTLs not identified in hepatocyte or bulk data and found 12 eQTLs in non-hepatocyte cell types. For example, LSECs represent 11% of liver cells (**Figure 1**), where they maintain inactivation of hepatic stellate cells (a mesenchymal cell subtype), exerting anti-fibrotic effects^65^. LSECs could also directly contribute to liver fibrosis by producing fibronectin and modulating extra cellular matrix metabolism^66^. We found an LSEC eQTL for *ADAMTS12*, a gene whose absence is associated with increased fibrosis in mice^67^. *ADAMTS12* is not identified as an eQTL in hepatocyte or bulk tissue analyses (**Figure 3**, **Table S8**), potentially due to the low proportion of LSECs in bulk liver tissue. This LSEC eQTL has a proxy variant (rs1037104, r^2^ = 0.99) located within an intronic accessible peak detected only in LSECs. This signal provides evidence of a genetic regulatory mechanism for *ADAMTS12* in LSECs. Thus, cell-type eQTLs can detect signals and elucidate regulatory mechanisms missed in less abundant cell types and bulk analyses.

**Figure 3:**
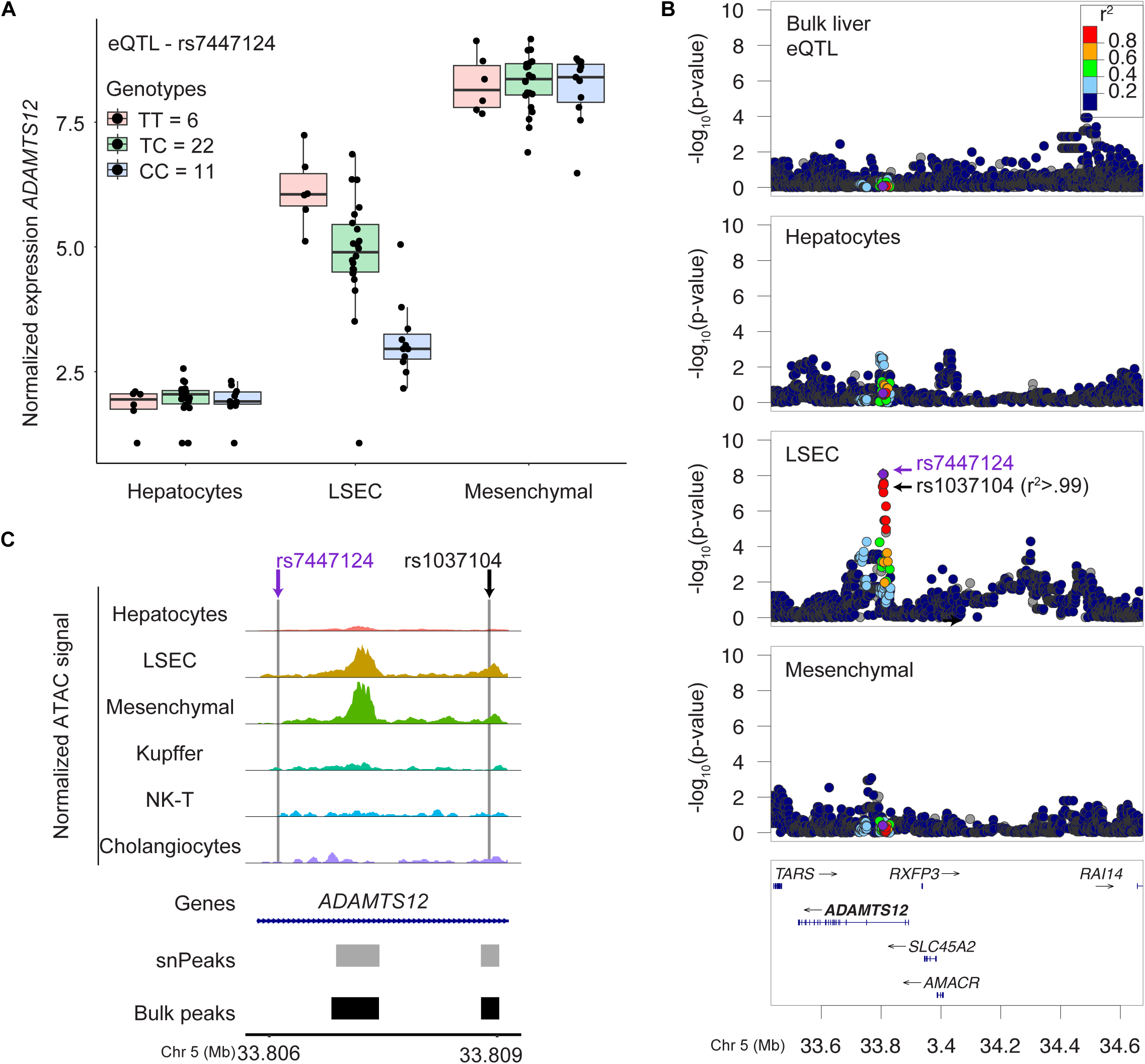
eQTL for *ADAMTS12* detected in LSECs. (**A**) rs7447124 is associated with *ADAMTS12* expression in LSECs but not in hepatocytes nor mesenchymal cells. Box plots show normalized expression levels by genotypes, with individual points representing samples. (**B**) Variant association with *ADAMTS12* in bulk liver from liver GTEx v8 and by cell type. Variants are colored by linkage disequilibrium with lead variant rs7447124 (purple dot). rs1037104 (black arrow) is in high LD with the lead variant (r^2^>.99). (**C**) rs1037104 is located in an intronic *ADAMTS12* accessible chromatin peak that only passed the detection threshold in LSEC. Gray bars below indicate regions defined as peaks in one or more cell types and black bars indicate regions defined as peaks in bulk liver tissue.

To link genetically regulated peaks to genetically regulated genes, we identified cell-type caQTL signals shared with bulk liver eQTLs. We identified caQTLs for 323 hepatocyte caPeaks that were shared (r^2^ ≥ 0.5) with bulk liver eQTLs for 344 genes; some regulatory elements may influence the expression of more than one gene (**Table S9**). 78% of the shared caQTL – eQTL signals showed alleles associated with both increased chromatin accessibility and increased gene expression, suggesting that these regulatory elements primarily function as promoters or enhancers. Conversely, the 22% of shared signals exhibiting opposite effects may represent transcriptional silencers or repressive elements in the hepatocyte regulatory landscape, a pattern that aligns with previous studies^11,13,68^. These findings demonstrate that cell-type caQTLs uncover both activating and repressive mechanisms by which genetic variants regulate gene expression.

### Predicting cell-type context for disease associations

To determine if genetic risk for certain disease-relevant traits is concentrated in specific liver cell types, we tested for heritability enrichment of GWAS traits in cell-type accessible chromatin peaks using stratified LD score regression. We found that loci associated with plasma levels of liver enzymes were enriched (FDR < 0.05) in hepatocytes, LSEC, cholangiocytes, and Kupffer cells (**Figure S10**, **Table S10**). Hepatocytes were enriched in loci for diagnosed diabetes and proteins produced by the liver such as insulin-like growth factor 1. Kupffer and NK-T cells were enriched in loci for counts of leukocytes and other immune cell types, consistent with their role in immunity. Mesenchymal cells were enriched in loci for blood pressure and eye measurement traits, potentially related to the role of hepatic stellate cells in vitamin A homeostasis^69^. These results suggest that accessible chromatin regions in different cell types are important for different traits.

To connect regulatory elements to GWAS signals, we identified shared signals between cell-type caQTLs and GWAS signals for 17 cardiometabolic and liver-relevant traits. Among 1,880 hepatocyte caQTLs, 205 were shared with at least 1 GWAS signal (r^2^ ≥ 0.5) (**Table 2, Table S9**). For example, 13% of GWAS signals for plasma levels of gamma-glutamyl transferase (GGT), an enzyme whose elevated levels are a marker of liver damage, and 8% of GWAS signals for low-density lipoprotein (LDL) cholesterol levels were shared with hepatocyte caQTLs, consistent with the role of hepatocytes in liver enzyme production and cholesterol metabolism. In a parallel analysis of eQTLs, we identified 10 eQTL signals shared with GWAS signals for liver enzymes, blood cholesterol, or triglyceride levels (r^2^ ≥ 0.5). These shared signals identify the chromatin accessible regions and genes that may be involved in genetic regulation of cardiometabolic diseases.

**Table 2:** GWAS signals shared with hepatocyte caQTL.

| **Trait** | **Total GWAS**<br>**signals analyzed** | **GWAS signals**<br>**shared with**<br>**caQTLs (%)** |
| --- | --- | --- |
| Gamma-Glutamyl Transferase<sup>3</sup> | 200 | 26 (13%) |
| Alanine Transaminase<sup>3</sup> | 149 | 16 (11%) |
| Alkaline Phosphatase<sup>3</sup> | 288 | 27 (9%) |
| 2-Hour Glucose<sup>4</sup> | 12 | 1 (8%) |
| Low-Density Lipoprotein<sup>1</sup> | 687 | 53 (8%) |
| Total Cholesterol<sup>1</sup> | 828 | 59 (7%) |
| C-Reactive Protein<sup>59</sup> | 438 | 27 (6%) |
| Triglycerides<sup>1</sup> | 705 | 37 (5%) |
| Vitamin D<sup>58</sup> | 142 | 7 (5%) |
| Body Mass Index<sup>57</sup> | 940 | 40 (4%) |
| High-Density Lipoprotein<sup>1</sup> | 814 | 35 (4%) |
| Hemoglobin A1c<sup>4</sup> | 96 | 3 (3%) |
| Type 2 Diabetes<sup>5</sup> | 402 | 12 (3%) |
| Coronary Artery Disease<sup>2</sup> | 241 | 6 (2%) |
| Fasting Glucose<sup>4</sup> | 96 | 2 (2%) |
| Waist-to-Hip Ratio Adjusted for BMI<sup>6</sup> | 463 | 11 (2%) |

Shared signals are the number of GWAS signals for which the lead variant had moderate LD ( $r^2 > 0.5$ ) with the lead variant of a hepatocyte caQTL.

To identify cell-type eQTL and caQTL signals shared with a broader set of GWAS traits, we evaluated signals from the GWAS catalog^60^. We found 399 cell-type caQTLs shared with GWAS signals for over 2000 traits (r^2^ > 0.5). Similarly, we found 25 cell-type eQTLs shared with ∼200 traits (r^2^ > 0.5, **Table S11**). For example, we identified an eQTL in Kupffer cells for *ITGAD* that shared its lead variant with a GWAS signal for platelet^70^ and reticulocyte^71^ counts (**Figure 4**). *ITGAD* encodes an alpha integrin that forms part of the integrin receptors expressed on Kupffer cells^72^. The eQTL lead variant resided in the *ITGAD* promoter region, which was most accessible in Kupffer cells. The rs8050500-C allele was associated with increased *ITGAD* expression and higher platelet and reticulocyte counts. This signal was also present in bulk liver eQTL for *ITGAD*. This example and others (**Table S11**) show that cell-type eQTLs and caQTLs for less abundant cell types can inform mechanisms for a wide range of complex traits.

**Figure 4:**
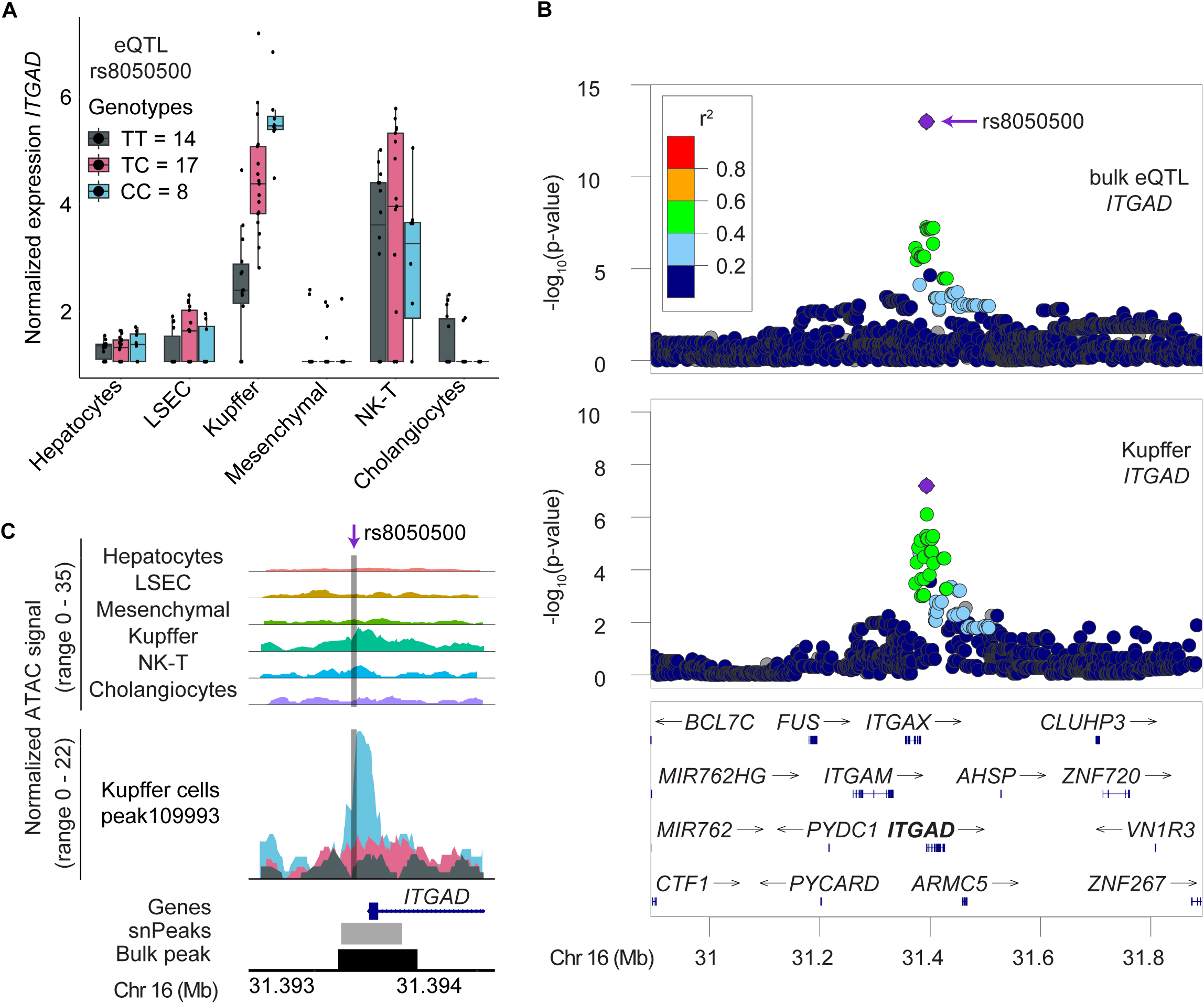
Cell-type eQTL signal shared with a GWAS signal for platelet and reticulocyte counts. (**A**) rs8050500 is associated with *ITGAD* expression only in Kupffer cells. Box plots show normalized expression levels by genotypes, with individual points representing samples. (**B**) Shared eQTL signal for *ITGAD* detected in bulk liver (GTEx) and in Kupffer cells. Lead variant rs8050500 (purple dot) in both signals is also associated with platelet^70^ and reticulocyte^71^ counts (not shown). (**C**) Coverage plot showing normalized ATAC-seq signal across cell types (top) and in an expanded view of Kupffer cells stratified by genotype rs8050500 (bottom). At the *ITGAD* promoter, individuals with rs8050500-CC genotype showed more accessible chromatin in Kupffer cells than the TC or TT genotypes. Colors correspond to genotypes shown in (A).

GWAS signals shared with both cell-type eQTLs and caQTLs reveal regulatory elements potentially regulating the expression of genes involved in cardiometabolic traits. At the *XKR9* locus, we identified a hepatocyte caQTL shared with a hepatocyte eQTL for *XKR9* (r^2^ = 0.97) and a GWAS signal for triglycerides (r^2^ = 0.87, **Figure 5, Table S12**); these signals were also observed in bulk liver QTL studies^10,11^ (**Figure S11**). The caQTL proxy variant rs6472538 (r^2^ = 0.99) resides within 30 bp of the caPeak center, and the rs6472538-A allele was associated with increased chromatin accessibility, increased *XKR9* expression, and elevated triglyceride levels. *XKR9* functions as a phospholipid scramblase that promotes phosphatidylserine exposure on the surface of apoptotic cells^73^. These shared signals provide a putative functional variant, altered regulatory element, target gene, and cell type of action for this triglycerides GWAS signal. We additionally found 80 hepatocyte caQTLs that were shared with bulk liver eQTLs for 87 genes and a GWAS signal for cardiometabolic traits (r^2^ ≥ 0.5, **Table S9**). This integrative approach demonstrates that combining cell-type chromatin accessibility signals with cell-type and bulk gene expression signals can help fine-map potential functional variants, regulatory regions, and disease-associated genes.

**Figure 5:**
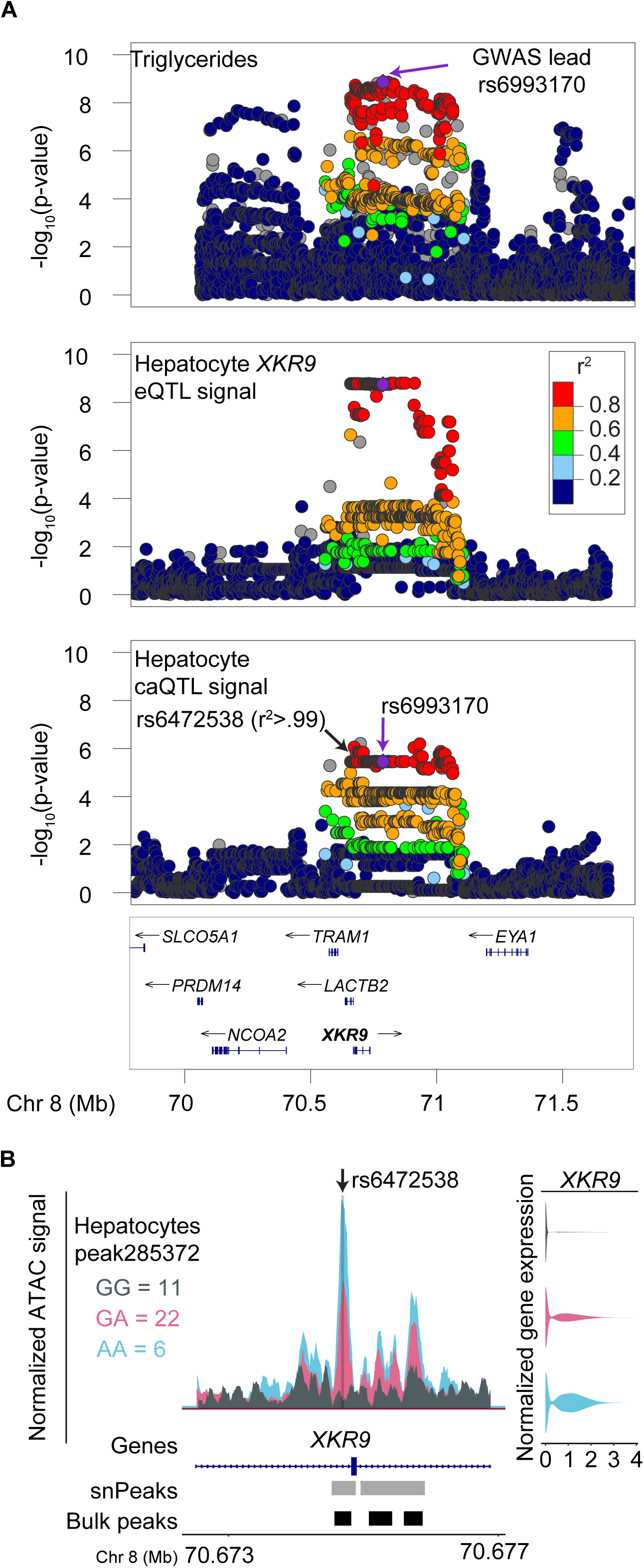
Cell-type caQTL signal shared with a cell-type eQTL signal and a GWAS signal for triglycerides. (**A**) A GWAS signal for triglycerides (lead variant rs6993170) is shared with a hepatocyte eQTL for *XKR9* and a hepatocyte caQTL. One LD proxy variant for rs6993170 is rs6472538 (r^2^>.99). (**B**) Coverage plot showing normalized ATAC-seq signal in hepatocytes stratified by genotype of rs6472538, located in an *XKR9* intron. Individuals with the rs6472538-AA genotype showed more hepatocyte accessible chromatin and higher hepatocyte gene expression than individuals with the GA or GG genotypes. Gray bars below indicate regions defined as peaks in hepatocytes and black bars indicate regions defined as peaks in bulk liver tissue.

### Cell-type annotation of bulk liver accessible chromatin

A key advantage of cell-type resolution of chromatin profiles is the ability to annotate cell types for caQTL detected in bulk tissue analyses. Using cell-type data, we were able to annotate cell types for 87% (30,751) of the bulk caPeaks^11^ (**Supplemental Data 1**). While the majority of bulk caPeaks (62%) were detected across multiple cell types, we detected 3,112 bulk caPeaks in only a single non-hepatocyte cell type (**Figure 6**, **Table S13**). This finding underscores the value of cell-type annotation for bulk caPeaks, enabling regulatory signals to be predicted in less abundant cell types even without identifying cell-type QTL.

**Figure 6:**
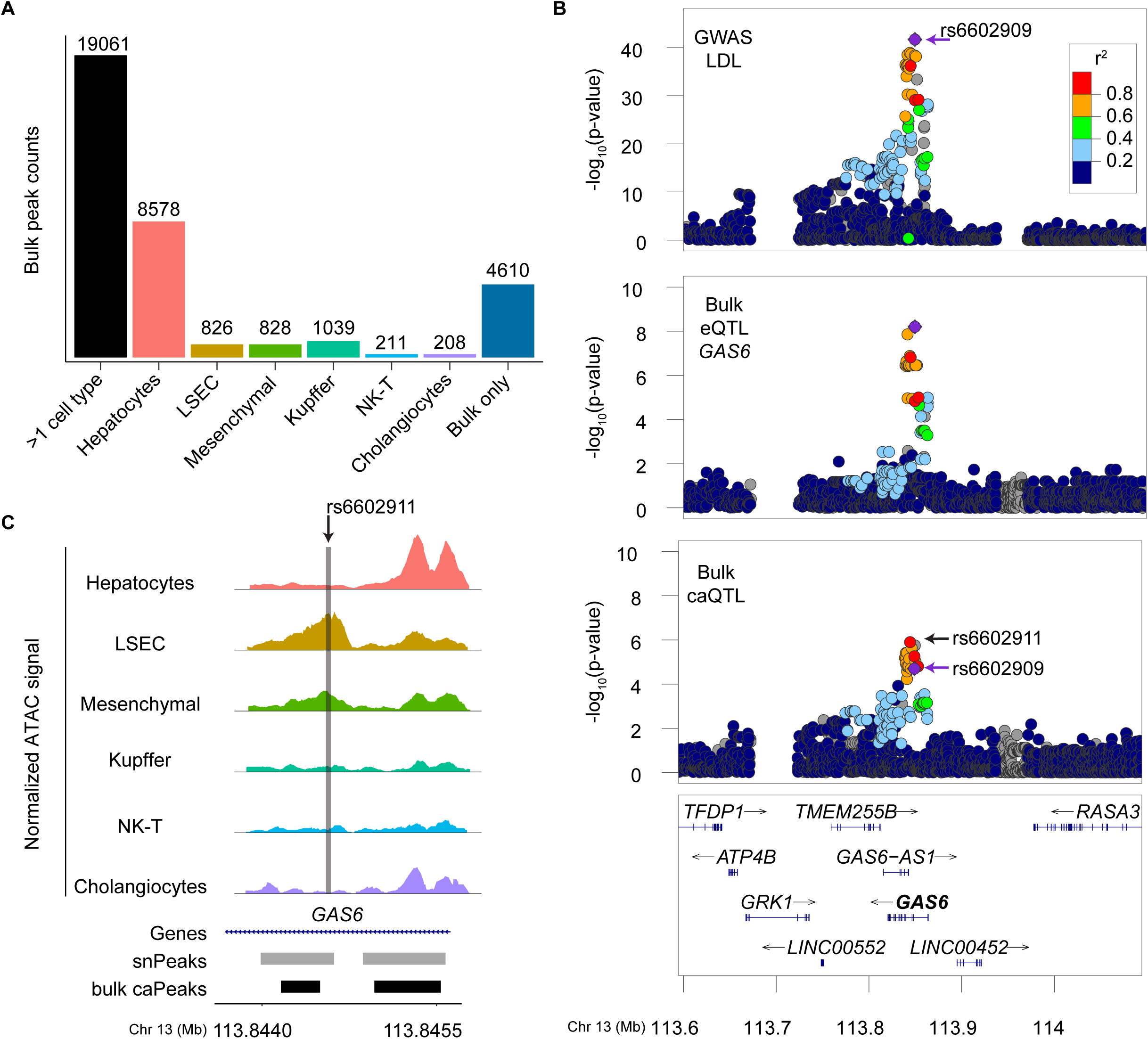
Liver cell types that correspond to bulk caQTLs and a subset that are colocalized with one or more GWAS signals. (**A**) Distribution of bulk caQTL peaks based on the cell types in which the same region of chromatin was accessible. Most peaks were detected in multiple cell types (19,061) or only in hepatocytes (8,578). 3,112 caQTLs were only detected in less abundant cell types. A subset (917) of these 30,751 bulk caQTL peaks colocalized with cardiometabolic GWAS signals. (**B**) GWAS signal for LDL with lead variant rs6602909 is colocalized with a bulk eQTL for *GAS6* and a bulk caQTL conditioned on rs74118417 (**methods**). One LD proxy variant for rs6602909 is rs6602911 (r^2^=.82). (**C**) The bulk caQTL lead variant rs6602911 overlaps accessible chromatin in LSEC and mesenchymal cells. The coverage plot shows normalized ATAC-seq signal at the *GAS6* locus for each cell type.

Within complex tissues such as the liver, each cell type contributes differently to physiological homeostasis and disease pathogenesis. Mapping GWAS loci to regulatory elements of these functionally distinct cell types helps uncover relevant mechanisms for liver diseases. Thus, we annotated the 998 bulk caQTLs^11^ that colocalized with GWAS signals of cardiometabolic traits by overlapping these bulk caPeaks with cell-type peaks. We identified putative cell type(s) for 917 of these bulk caPeaks and the distribution of these colocalized bulk caQTLs across different cell types corresponded with cell-type proportions in our dataset (**Table S14**). One example is for a bulk caQTL for a peak detected only in LSEC and mesenchymal cells and that colocalized with GWAS signals for alkaline phosphatase (ALP), total and low-density lipoprotein (LDL) cholesterol, and triglyceride levels (**Figure 6**). The bulk caPeak lies in the intron of *GAS6,* and its lead variant also colocalized with a bulk liver eQTL for the same gene. *GAS6* has been implicated in hepatic stellate cell activation^74^, consistent with the role of mesenchymal cells in fibrosis. Overall, these analyses highlight the importance of cell-type resolution in nominating cell types for signals identified in bulk analyses, particularly for less abundant cell populations

## Discussion

We profiled the human liver at single-nucleus resolution and revealed cell-type regulatory landscapes relevant to liver-related and cardiometabolic traits. Through multiome analysis of snRNA-seq and snATAC-seq data from 39 human liver samples, we identified distinct regulatory profiles across six major cell types, detecting 1,885 caQTLs and 67 eQTLs. This cell-type resolution improves our understanding of liver genetics by suggesting the cellular context in which disease-associated variants operate. We showed that a single nucleus approach can help dissect how genetic variants contribute to various physiological and pathological conditions such as metabolic disorders in hepatocytes and inflammatory responses in Kupffer cells. By linking cell-type regulatory elements to GWAS signals and annotating previously identified bulk tissue associations, we can predict regulatory mechanisms underlying disease associations that are obscured in conventional bulk tissue analyses.

Our cell-type resolved QTL analyses suggested the cellular origins of regulatory mechanisms underlying several GWAS associations for liver-related traits. We identified an eQTL for *ITGAD* in Kupffer cells, which showed that genetic variation may influence integrin-mediated leukocyte adhesion in liver-resident macrophages. Integrins regulate Kupffer cell interactions with circulating immune cells and platelets, directly affecting their function in platelet clearance^80^. Additionally, these receptors participates in activation and expansion of Kupffer cells in non-alcoholic steatohepatitis^72^. The shared genetic architecture between these cell-type eQTLs and GWAS signals enhances our understanding of GWAS loci by providing biologically interpretable mechanisms to statistical associations, pinpointing the cellular origins of these regulatory effects.

Our cell-type approach demonstrates value for annotating regulatory elements identified in bulk liver studies. Despite a modest size of 39 samples, we successfully detected 70% of the accessible chromatin regions detected in a bulk liver chromatin accessibility study of 138 samples^11^, while identifying 70,884 cell-type peaks undetected in bulk data. Most of these novel peaks were detected only in non-hepatocyte cell types, highlighting that single-nucleus profiling effectively captures regulatory elements masked by cellular heterogeneity in bulk analyses. By annotating cell types for 30,751 previously reported bulk liver caQTLs^11^, we enhanced the interpretation of regulatory signals, including a subset of 917 regulatory elements at GWAS signals, providing cellular context for disease-relevant genetic variation. For example, we annotated the accessibility of a bulk caPeak to LSEC and mesenchymal cells located within an intron of *GAS6*. This bulk caQTL also colocalized with an eQTL for the same gene, providing mechanistic insight into how genetic variation might influence fibrotic processes in a cell-type manner. *GAS6* encodes growth arrest-specific 6, a signaling hub of liver fibrosis^75^, and mice deficient in *Gas6* show less susceptibility to steatosis and fibrosis^74^. *GAS6* is involved in cellular differentiation, immune response, and the activation of hepatic stellate cells^74^. The association of this signal with elevated alkaline phosphatase levels and other cardiometabolic traits underscores its importance in metabolic liver diseases and may offer a promising target for anti-fibrotic therapies.

While this study demonstrates the value of cell-type resolved regulatory analysis in liver tissue, several limitations shape opportunities for future research. Due to statistical power constraints, we cannot definitively establish cell-type specificity of quantitative trait loci, as the absence of detected signals in less abundant cell populations may reflect insufficient detection sensitivity rather than true biological absence of regulatory effects. Our attempts to identify additional eQTLs by lowering UMI thresholds during nuclei filtration produced fewer significant results, likely due to inclusion of non-nucleus barcodes or contaminated nuclei (data not shown). Future studies including more individuals with defined disease states would enable better identification of context-dependent regulatory effects. Finally, integrating emerging technologies like cell-type Hi-C with single-nucleus ATAC and RNA sequencing offers promising approaches for connecting regulatory elements to target genes through direct chromatin contact.

Despite these limitations, our findings demonstrate the utility of single-nucleus approaches for charactering the genetic variation in liver cell types. Coupling single-nucleus and bulk profiling enhanced our ability to dissect the cellular contexts of genetic regulation in the liver and identified cell-type regulatory elements relevant to cardiometabolic disease.

## Acknowledgments

This study was supported by NIH grants R01DK072193, UM1DK126185 and T32HL069768 (HJP). We thank the donors of liver samples, which were obtained from the Liver Tissue Cell Distribution System supported by NIH contract N01DK70004/ HHSN267200700004C. We thank members of the Mohlke lab for helpful discussions on computational analyses.

We gratefully acknowledge the technical support from UNC Cores. The UNC High Throughput Sequencing Facility is supported by the University Cancer Research Fund, Comprehensive Cancer Center Core Support grant (P30-CA016086) and the UNC Center for Mental Health and Susceptibility grant (P30-ES010126). The Advanced Analytics Core is funded through the UNC Center for Gastrointestinal Biology and Disease grant (P30 DK034987). The UNC Pathology Services Core Facility is supported in part by an NCI Center Core Support Grant (P30-CA016086).

We also thank GTEx for liver tissue gene expression data. The Genotype-Tissue Expression (GTEx) Project was supported by the Common Fund of the Office of the Director of the National Institutes of Health, and by NCI, NHGRI, NHLBI, NIDA, NIMH, and NINDS. The data used for the analyses described in this manuscript were obtained from the GTEx Portal.

## Data and Code Availability

All raw and processed genotype, snATAC-seq, and snRNA-seq data will be made available in GEO upon publication.

---

**Figure S1:**
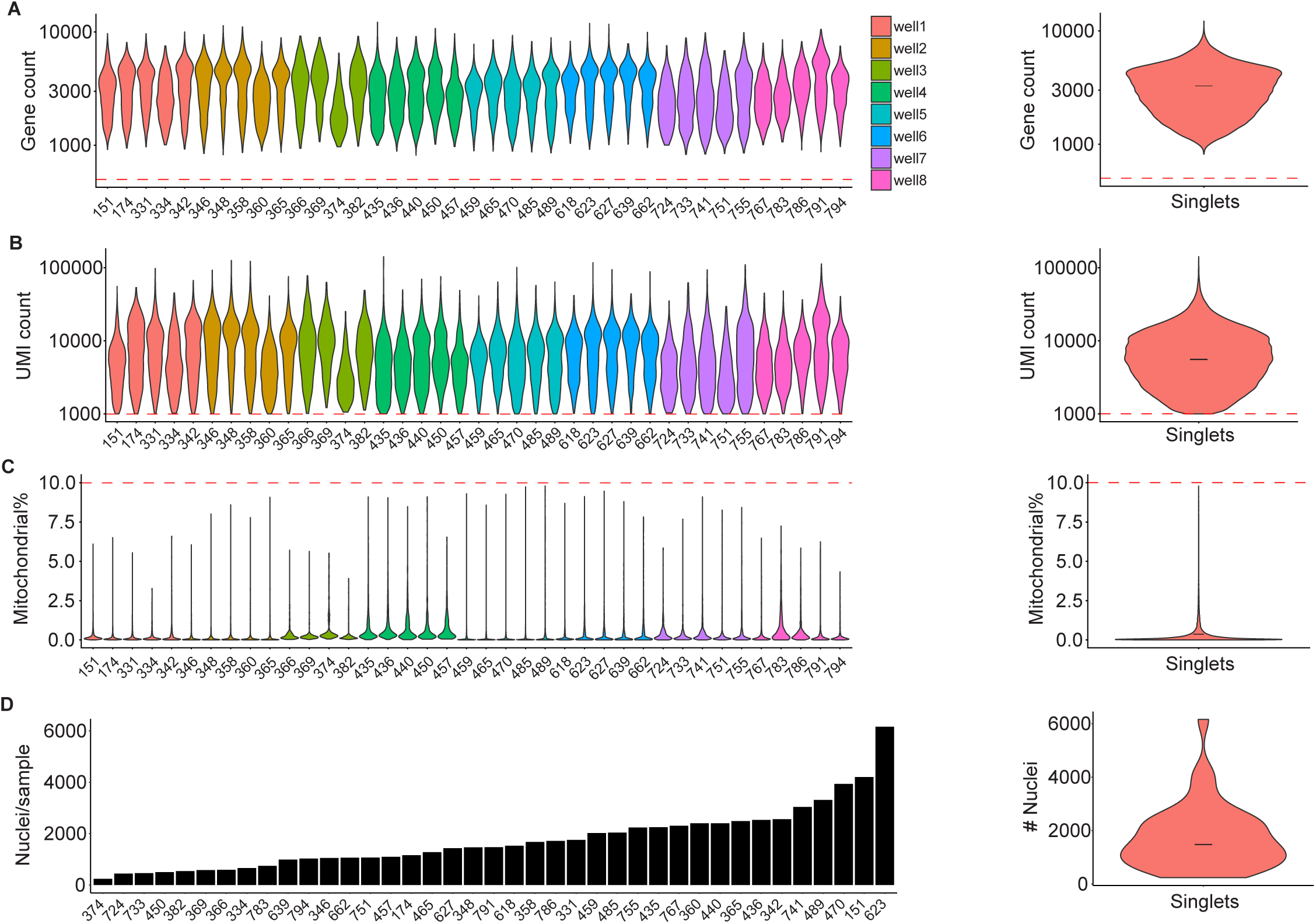
Quality of single-nucleus RNA sequencing data. Data for each metric is shown by processing batch (wells 1-8, left panels) and aggregated across all nuclei (right panels). (A) Gene count distribution per nucleus. (B) Unique molecular identifier (UMI) count distribution per nucleus. (C) Mitochondrial RNA content as percentage of total counts per nucleus. (D) Post-quality control nucleus count per donor sample and distribution across donors. n = 39 samples. Quality control thresholds are indicated by red dashed lines.

**Figure S2:**
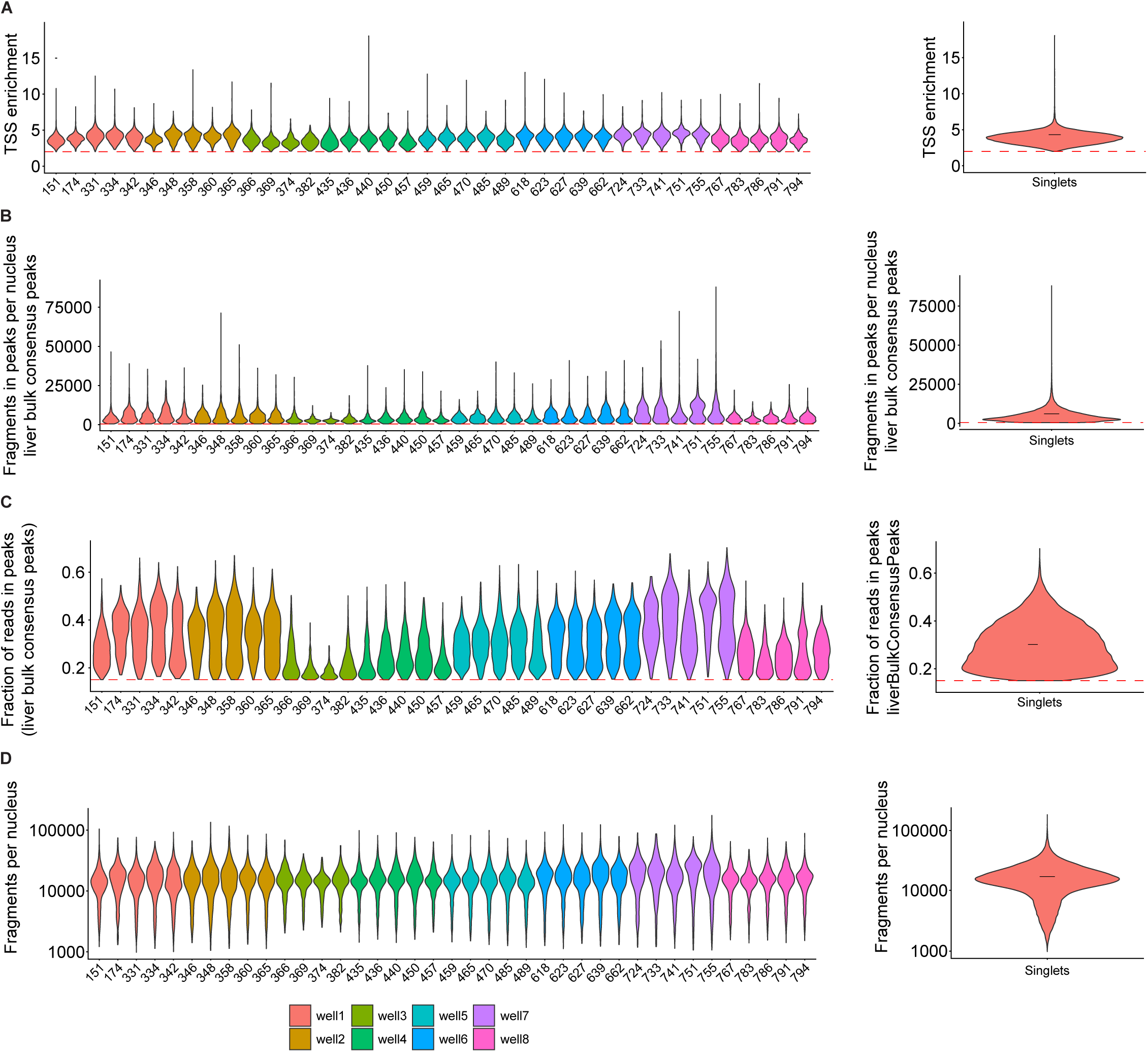
Quality of single-nucleus ATAC sequencing data. Data for each metric is shown by processing batch (wells 1-8, left panels) and aggregated across all nuclei (right panels). (A) Transcription start site (TSS) enrichment scores per nucleus calculated using EnsDb.Hsapiens.v86. (B) Number of fragments in peaks per nucleus. (C) Fraction of fragments in peaks (FRiP) per nucleus. Thresholds in panels (B and C) were applied to peaks with coordinates from bulk liver tissue. (D) Total fragment count distribution per nucleus. n = 39 samples. Quality control thresholds are indicated by red dashed lines.

**Figure S3:**
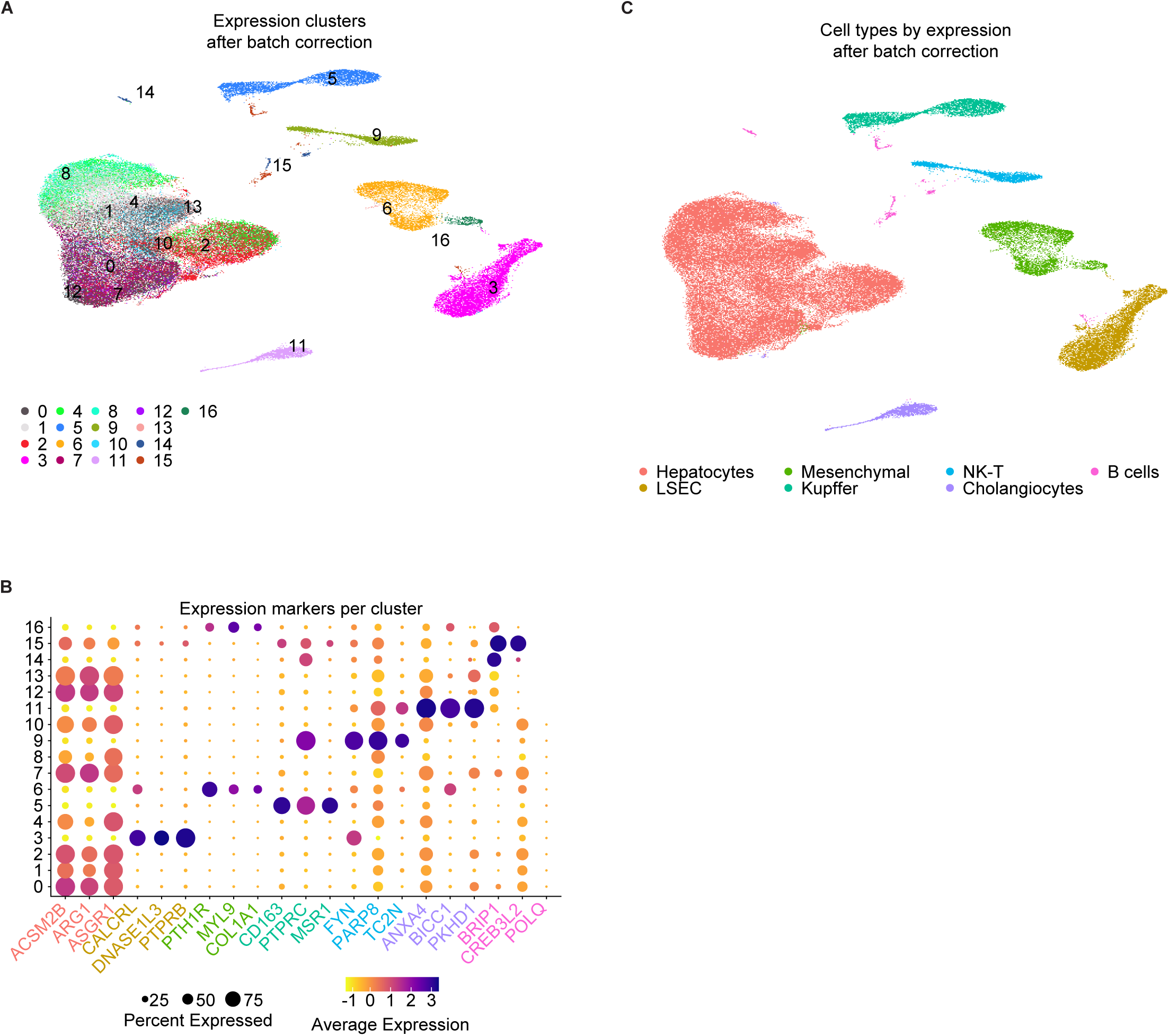
Cell type identification using single-nucleus RNA-seq data. (A) Initial clustering of expression data after batch correction showing 17 distinct clusters. (B) Established cell-type marker gene expression across identified clusters. Dot size represents the percentage of cells expressing the marker and color intensity shows the average expression level. (C) Cell-type annotations after batch effect correction of snRNA-seq data.

**Figure S4:**
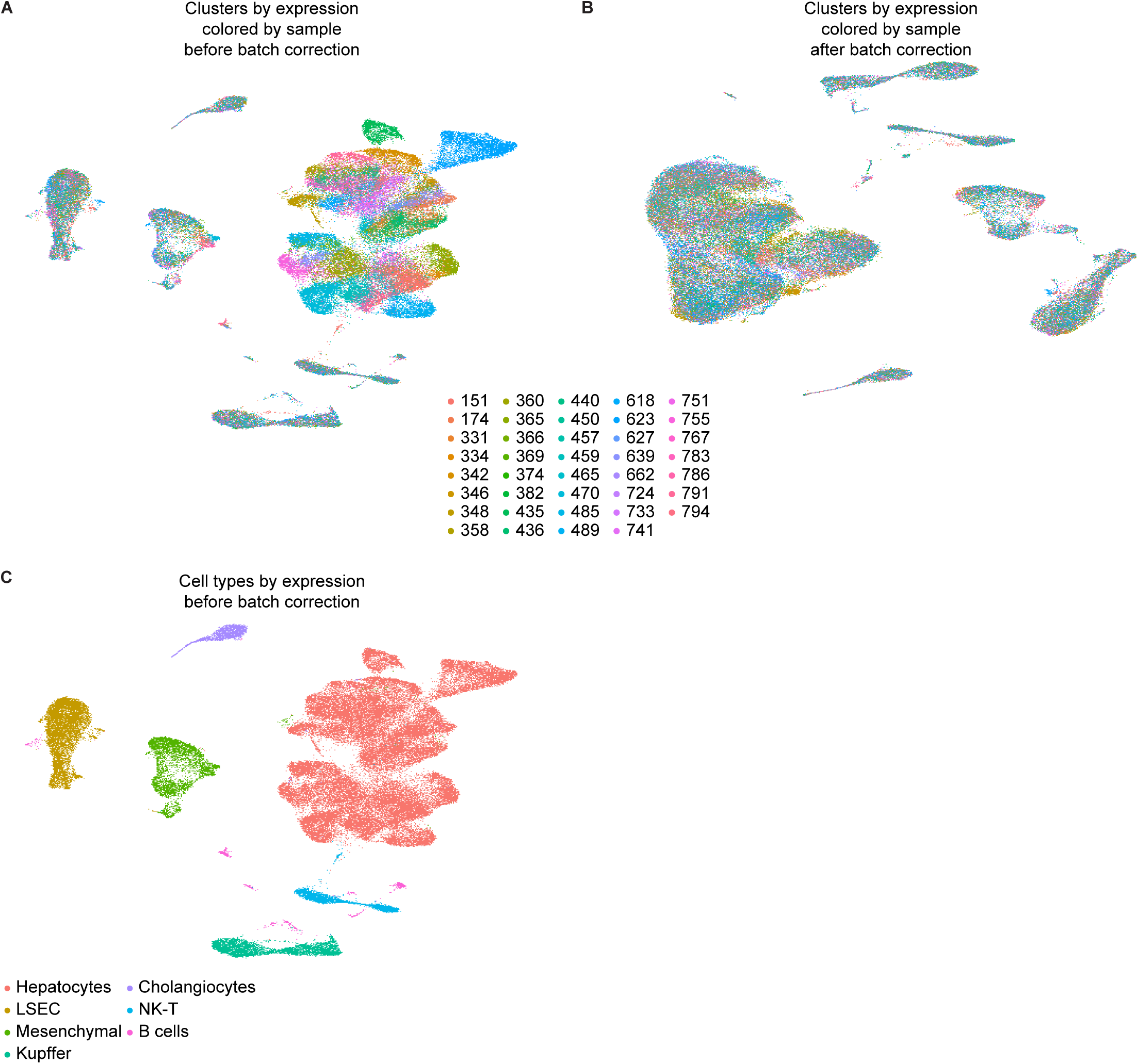
Batch correction in single-nucleus RNA-seq data. (A - B) Clusters of expression data before (A) and after (B) batch correction colored by sample. (C) Cell-type annotations prior to batch correction of RNA-seq data. Cell-type annotation after batch correction of RNA-seq data is shown in Figure S3.

**Figure S5:**
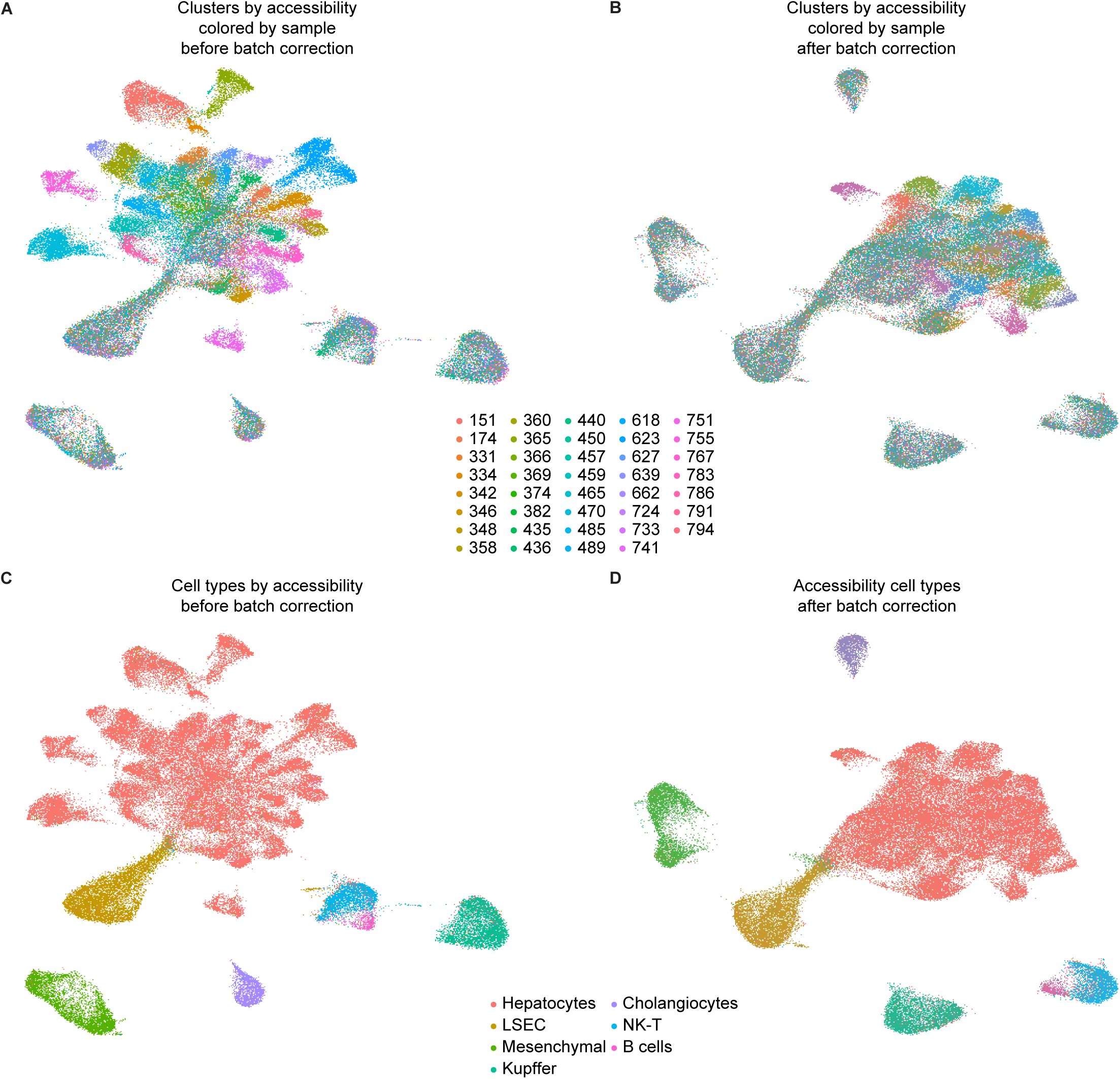
Batch correction in single-nucleus ATAC-seq data. (A - B) Clusters of expression data before (A) and after (B) batch correction colored by sample. (C - D) Cell-type annotations before (C) and after (D) batch correction of RNA-seq data. 20 harmonized LSI components were used to recalculate clusters and UMAP projections.

**Figure S6:**
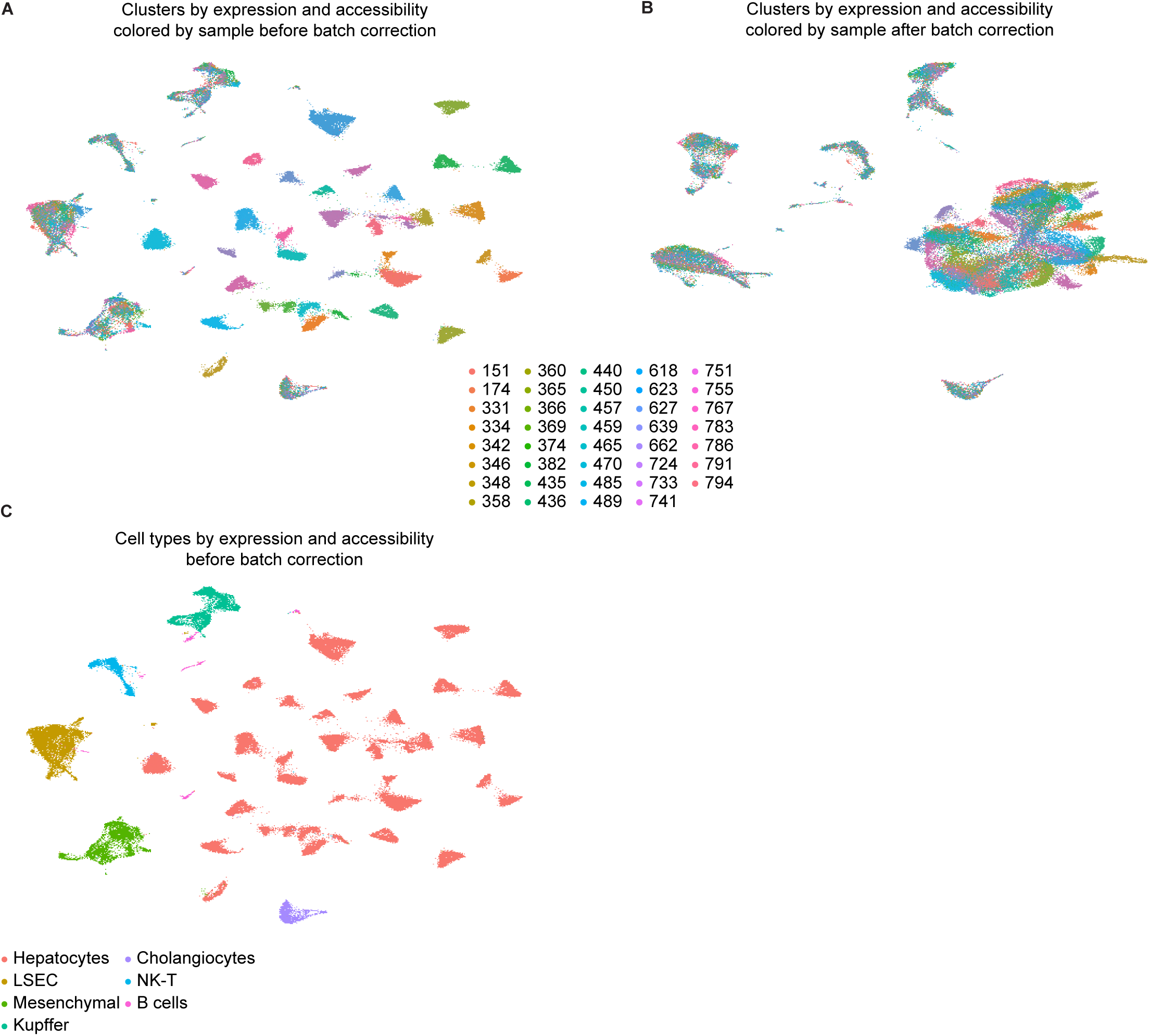
Batch correction in joint UMAP of single-nucleus RNA and ATAC data. (A - B) Clusters of joint profiles (snRNA-seq + snATAC-seq) before (A) and after (B) batch correction colored by sample. (C) Cell-type annotations of joint profiles before batch correction. Cell-type annotations of joint profules after batch correction is shown in Figure 1.

**Figure S7:**
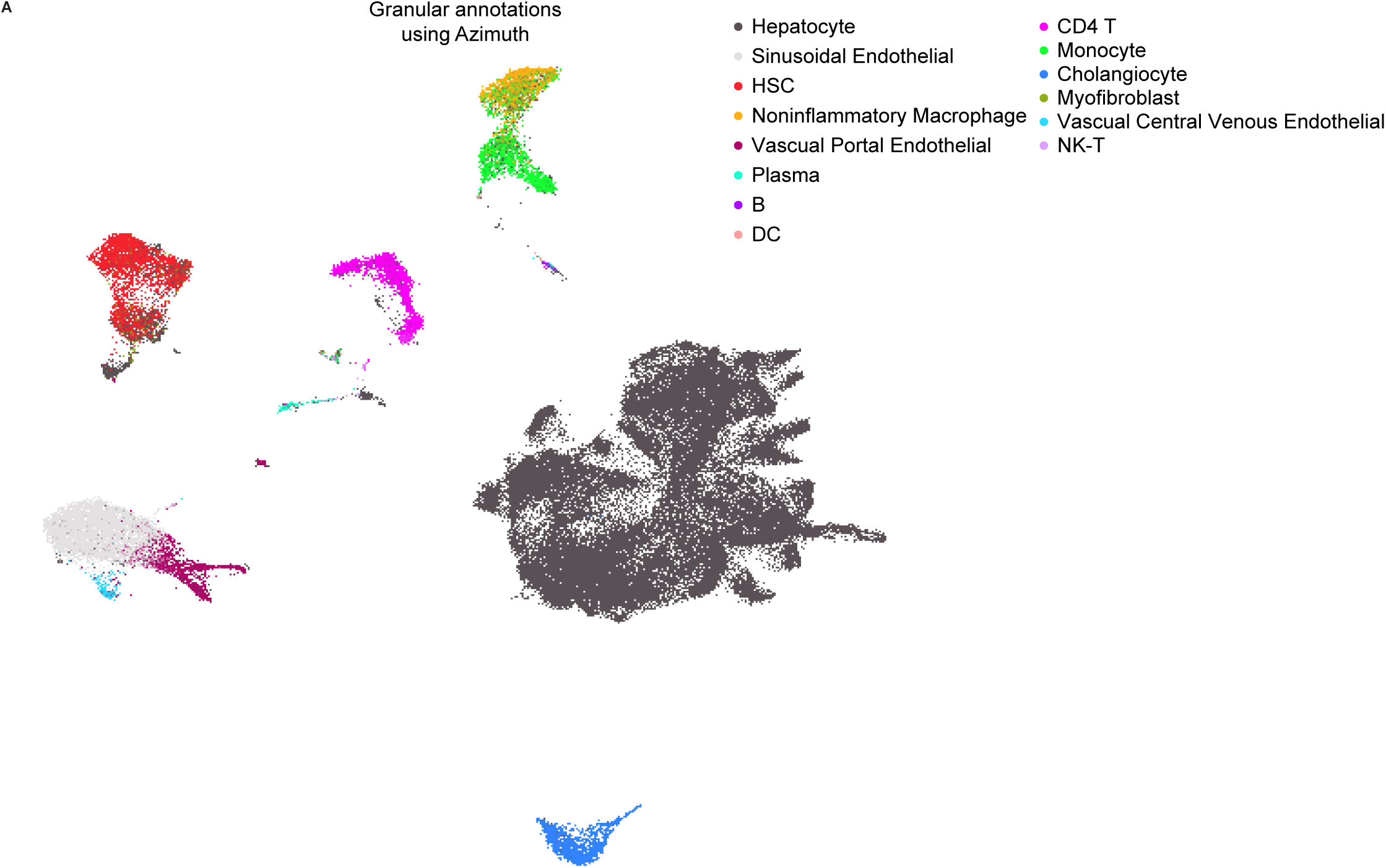
Granular cell-type annotations. (A) Detailed annotation of batch-corrected and snRNA+snATAC-seq integrated data using Azimuth reference mapping

**Figure S8:**
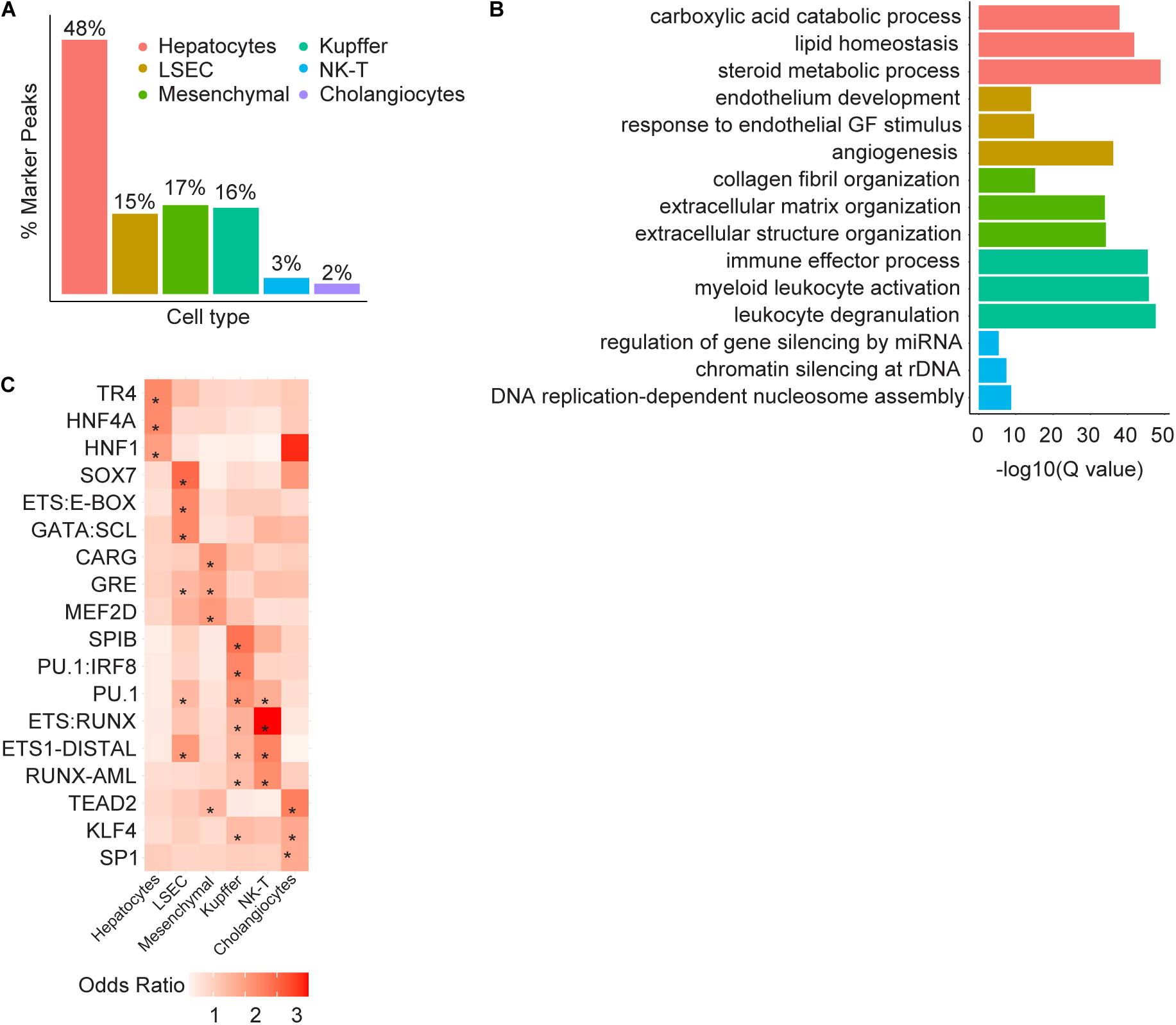
Cell-type marker peaks and their associated regulatory features. (A) Distribution of cell type marker peaks across major liver cell populations, ordered by cell-type proportion. Among 17,147 marker peaks, most are hepatocytes (48%), followed by mesenchymal cells (17%), Kupffer cells (16%), and LSECs (15%), while NK-T cells (3%) and cholangiocytes (2%) show fewer marker peaks. (B) Gene ontology terms of genes near marker peaks based on GREAT. The x-axis represents statistical significance as -log10(Q-value), and the three most significant terms per cell type are displayed. Bar colors denote cell types as in (A). (C) Heatmap showing transcription factor (TF) motif enrichment within marker peaks for each cell type, quantified by enrichment odds ratios (color intensity). Asterisks denote significant enrichment (FDR < 0.05). The three most enriched motifs per cell type are shown.

**Figure S9:**
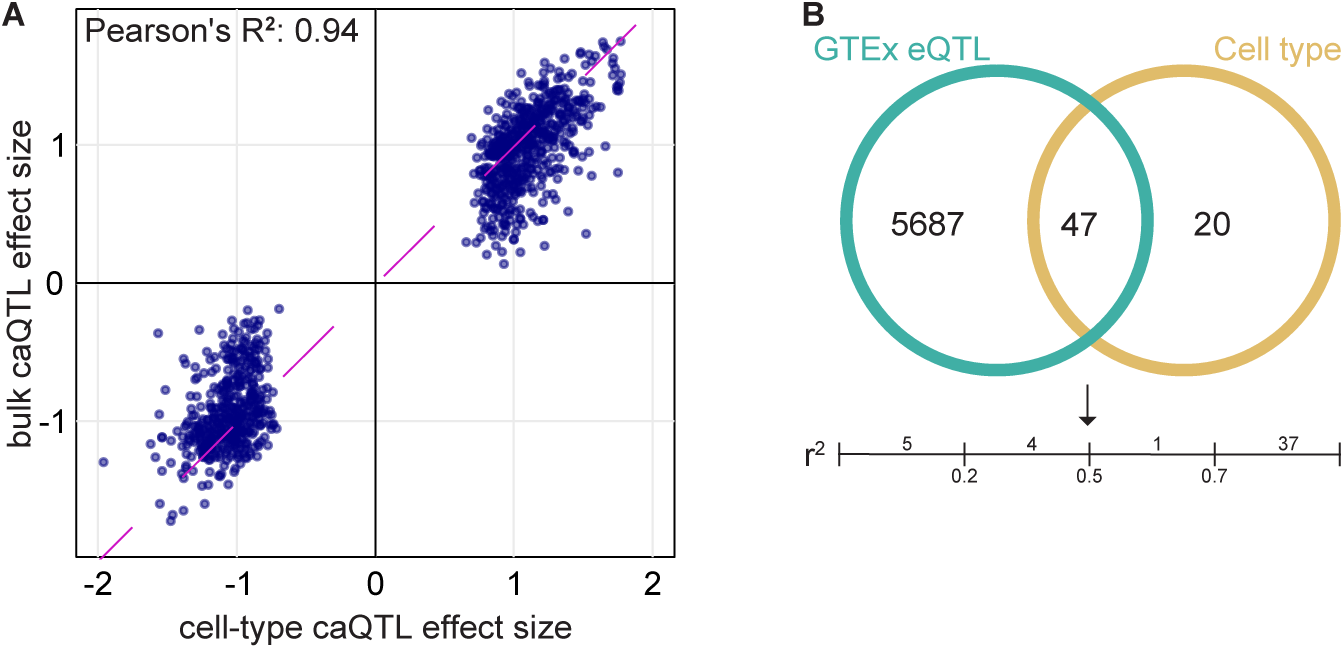
Comparison between bulk and cell-type QTLs. (A) The effect sizes (beta) of caQTLs identified in both the bulk liver tissue and one or more cell-type show strong concordance (Pearson’s R² = 0.94). Dots represents the 975 caQTLs that are lead variants both in bulk and cell-type data. (B) Venn diagram detailing the LD r^2^ between 47 genes with detected eQTLs both in GTEx bulk liver tissue (green) and our cell-type analysis (gold). We considered eQTLs with LD r^2^ >.5 as shared. While most eQTLs were only detected in the GTEx bulk analysis (5,687 genes), we identified 20 novel cell-type eQTLs for 20 genes. Among the 47 eQTLs detected in both datasets, the line plot below shows the LD r^2^ distribution between lead variants; 37 show strong linkage disequilibrium (r² ≥ 0.7).

**Figure S10:**
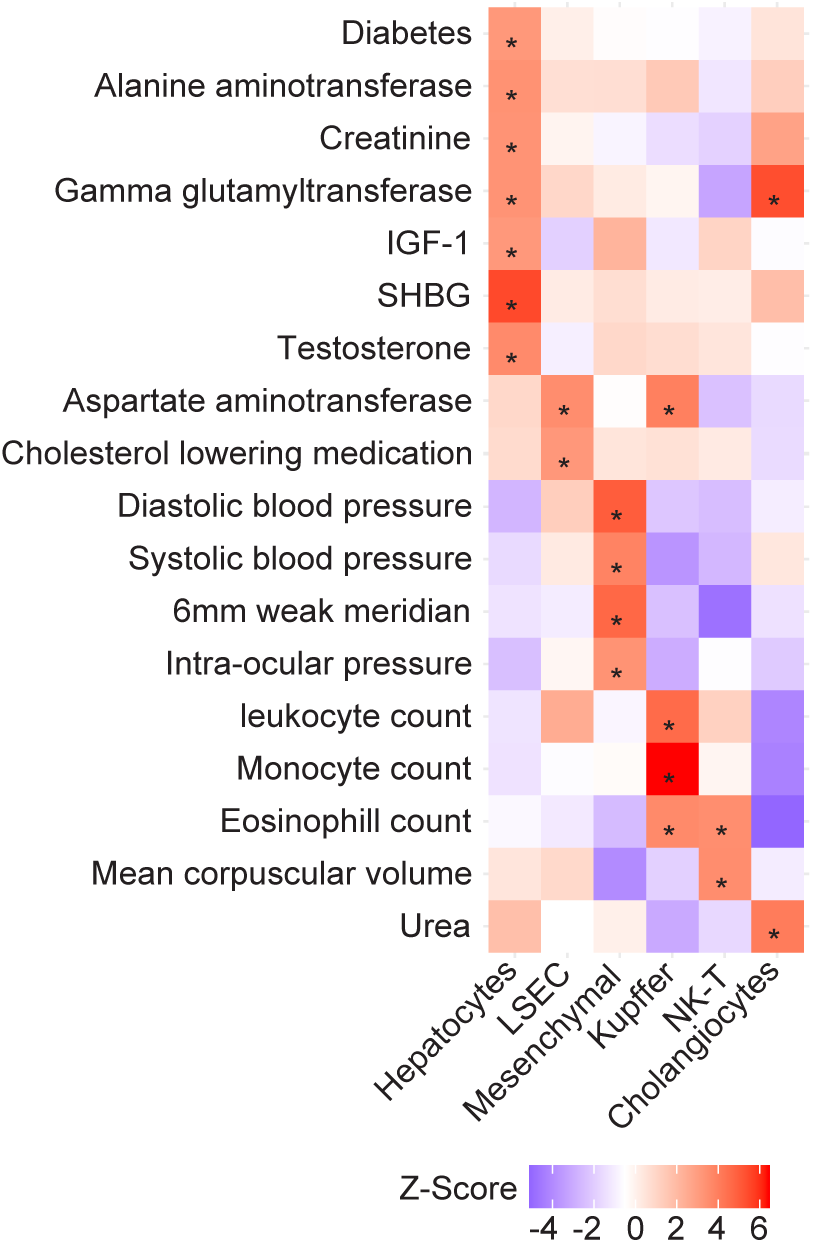
GWAS variants are enriched in cell-type peaks. Enrichment of GWAS signals in cell-type accessible chromatin regions. The heatmap shows z-scores of heritability enrichment across cell types. Asterisks indicate significant enrichments (FDR < 0.05).

**Figure S11:**
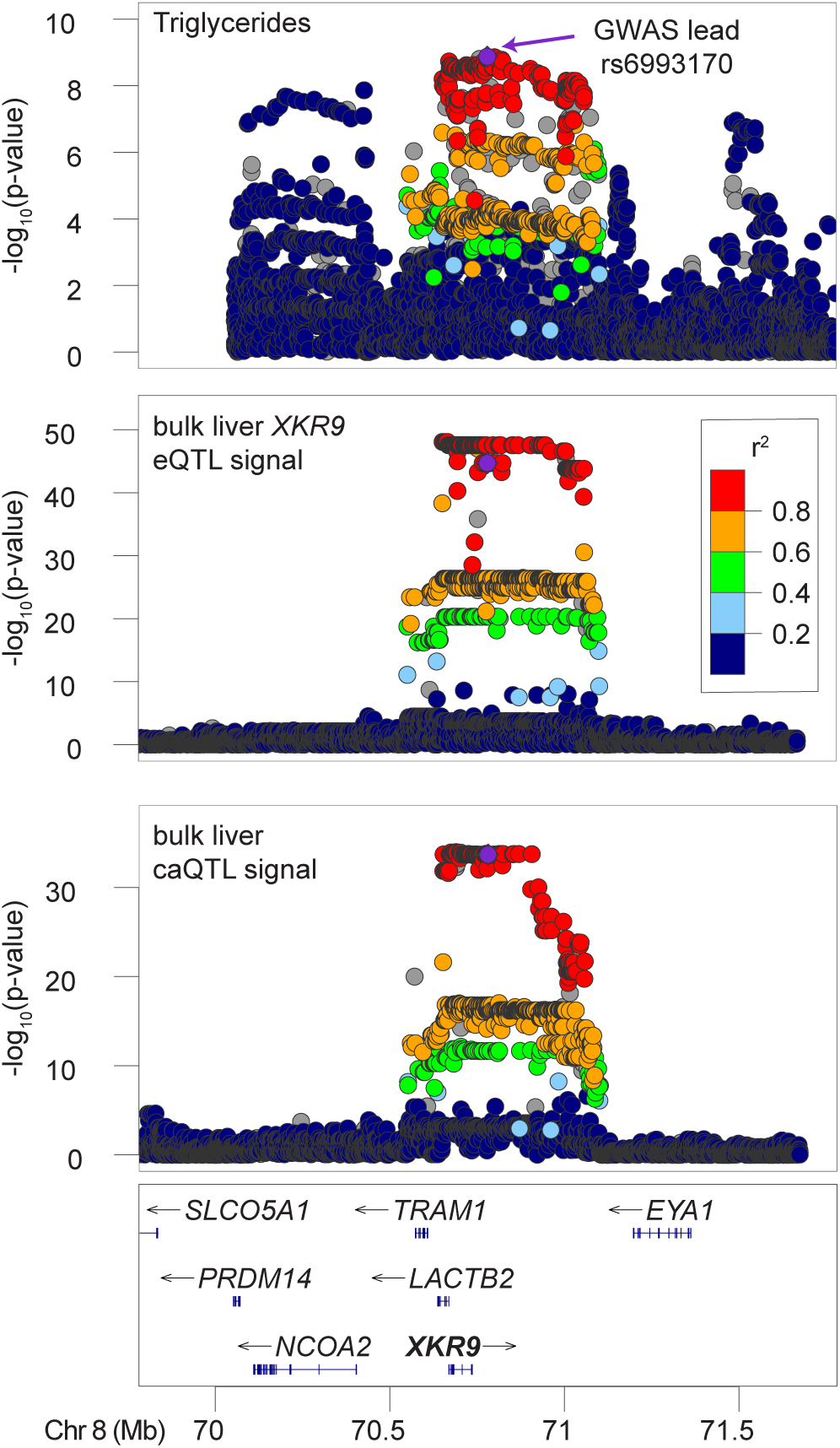
*XKR9* locus genetic variants identified in bulk datasets. A GWAS signal for triglycerides (lead variant rs6993170) is shared with a bulk liver eQTL for *XKR9* (rs13255886, r2=.99) and a bulk liver caQTL (rs12675271, r2=.99). The GWAS lead variant is shown by a purple dot in all the plots. The hepatocyte caQTL signal and the eQTL signal for *XKR9* are shown in Figure 5.

## References

1. Graham, S.E., Clarke, S.L., Wu, K.-H.H., Kanoni, S., Zajac, G.J.M., Ramdas, S., Surakka, I., Ntalla, I., Vedantam, S., Winkler, T.W., et al. (2021). The power of genetic diversity in genome-wide association studies of lipids. Nature 600, 675–679. 10.1038/s41586-021-04064-3.

2. Aragam, K.G., Jiang, T., Goel, A., Kanoni, S., Wolford, B.N., Atri, D.S., Weeks, E.M., Wang, M., Hindy, G., Zhou, W., et al. (2022). Discovery and systematic characterization of risk variants and genes for coronary artery disease in over a million participants. Nat Genet 54, 1803–1815. 10.1038/s41588-022-01233-6.

3. Pazoki, R., Vujkovic, M., Elliott, J., Evangelou, E., Gill, D., Ghanbari, M., van der Most, P.J., Pinto, R.C., Wielscher, M., Farlik, M., et al. (2021). Genetic analysis in European ancestry individuals identifies 517 loci associated with liver enzymes. Nat Commun 12, 2579. 10.1038/s41467-021-22338-2.

4. Chen, J., Spracklen, C.N., Marenne, G., Varshney, A., Corbin, L.J., Luan, J., Willems, S.M., Wu, Y., Zhang, X., Horikoshi, M., et al. (2021). The trans-ancestral genomic architecture of glycemic traits. Nat Genet 53, 840–860. 10.1038/s41588-021-00852-9.

5. Mahajan, A., Taliun, D., Thurner, M., Robertson, N.R., Torres, J.M., Rayner, N.W., Payne, A.J., Steinthorsdottir, V., Scott, R.A., Grarup, N., et al. (2018). Fine-mapping type 2 diabetes loci to single-variant resolution using high-density imputation and islet-specific epigenome maps. Nat. Genet. 50, 1505–1513. 10.1038/s41588-018-0241-6.

6. Pulit, S.L., Stoneman, C., Morris, A.P., Wood, A.R., Glastonbury, C.A., Tyrrell, J., Yengo, L., Ferreira, T., Marouli, E., Ji, Y., et al. (2019). Meta-analysis of genome-wide association studies for body fat distribution in 694 649 individuals of European ancestry. Human Molecular Genetics 28, 166–174. 10.1093/hmg/ddy327.

7. Absher, Devin (2012). An Integrated Encyclopedia of DNA Elements in the Human Genome. Nature 489, 57–74. 10.1038/nature11247.

8. Kundaje, A., Meuleman, W., Ernst, J., Bilenky, M., Yen, A., Heravi-Moussavi, A., Kheradpour, P., Zhang, Z., Wang, J., Ziller, M.J., et al. (2015). Integrative analysis of 111 reference human epigenomes. Nature 518, 317.

9. Finucane, H.K., Bulik-Sullivan, B., Gusev, A., Trynka, G., Reshef, Y., Loh, P.-R., Anttila, V., Xu, H., Zang, C., Farh, K., et al. (2015). Partitioning heritability by functional annotation using genome-wide association summary statistics. Nat Genet 47, 1228– 1235. 10.1038/ng.3404.

10. The GTEx Consortium (2020). The GTEx Consortium atlas of genetic regulatory effects across human tissues. Science 369, 1318–1330. 10.1126/science.aaz1776.

11. Currin, K.W., Perrin, H.J., Pandey, G.K., Alkhawaja, A.A., Vadlamudi, S., Musser, A.E., Etheridge, A.S., Broadaway, K.A., Rosen, J.D., Varshney, A., et al. (2025). Genetic effects on chromatin accessibility uncover mechanisms of liver gene regulation and quantitative traits. Genome Res, gr.279741.124. 10.1101/gr.279741.124.

12. Alasoo, K., Rodrigues, J., Mukhopadhyay, S., Knights, A.J., Mann, A.L., Kundu, K., Hale, and C., Dougan, G., and Gaffney, D.J. (2018). Shared genetic effects on chromatin and gene expression indicate a role for enhancer priming in immune response. Nat. Genet. 10.1038/s41588-018-0046-7.

13. Currin, K.W., Erdos, M.R., Narisu, N., Rai, V., Vadlamudi, S., Perrin, H.J., Idol, J.R., Yan, T., Albanus, R.D., Broadaway, K.A., et al. (2021). Genetic effects on liver chromatin accessibility identify disease regulatory variants. Am J Hum Genet 108, 1169–1189. 10.1016/j.ajhg.2021.05.001.

14. Liang, D., Elwell, A.L., Aygün, N., Krupa, O., Wolter, J.M., Kyere, F.A., Lafferty, M.J., Cheek, K.E., Courtney, K.P., Yusupova, M., et al. (2021). Cell-type-specific effects of genetic variation on chromatin accessibility during human neuronal differentiation. Nat Neurosci 24, 941–953. 10.1038/s41593-021-00858-w.

15. Calderon, D., Bhaskar, A., Knowles, D.A., Golan, D., Raj, T., Fu, A.Q., and Pritchard, J.K. (2017). Inferring Relevant Cell Types for Complex Traits by Using Single-Cell Gene Expression. Am J Hum Genet 101, 686–699. 10.1016/j.ajhg.2017.09.009.

16. Watanabe, K., Umićević Mirkov, M., de Leeuw, C.A., van den Heuvel, M.P., and Posthuma, D. (2019). Genetic mapping of cell type specificity for complex traits. Nat Commun 10, 3222. 10.1038/s41467-019-11181-1.

17. Zhang, K., Hocker, J.D., Miller, M., Hou, X., Chiou, J., Poirion, O.B., Qiu, Y., Li, Y.E., Gaulton, K.J., Wang, A., et al. (2021). A single-cell atlas of chromatin accessibility in the human genome. Cell 184, 5985–6001.e19. 10.1016/j.cell.2021.10.024.

18. Orchard, P., Manickam, N., Ventresca, C., Vadlamudi, S., Varshney, A., Rai, V., Kaplan, J., Lalancette, C., Mohlke, K.L., Gallagher, K., et al. (2021). Human and rat skeletal muscle single-nuclei multi-omic integrative analyses nominate causal cell types, regulatory elements, and SNPs for complex traits. Genome Res 31, 2258– 2275. 10.1101/gr.268482.120.

19. Chiou, J., Zeng, C., Cheng, Z., Han, J.Y., Schlichting, M., Miller, M., Mendez, R., Huang, S., Wang, J., Sui, Y., et al. (2021). Single-cell chromatin accessibility identifies pancreatic islet cell type- and state-specific regulatory programs of diabetes risk. Nat Genet 53, 455–466. 10.1038/s41588-021-00823-0.

20. Trefts, E., Gannon, M., and Wasserman, D.H. (2017). The liver. Curr. Biol. 27, R1147– R1151. 10.1016/j.cub.2017.09.019.

21. Almazroo, O.A., Miah, M.K., and Venkataramanan, R. (2017). Drug Metabolism in the Liver. Clin Liver Dis 21, 1–20. 10.1016/j.cld.2016.08.001.

22. Etheridge, A.S., Gallins, P.J., Jima, D., Broadaway, K.A., Ratain, M.J., Schuetz, E., Schadt, E., Schroder, A., Molony, C., Zhou, Y., et al. (2019). A New Liver Expression Quantitative Trait Locus Map From 1,183 Individuals Provides Evidence for Novel Expression Quantitative Trait Loci of Drug Response, Metabolic, and Sex-Biased Phenotypes. Clin. Pharmacol. Ther. 10.1002/cpt.1751.

23. Broadaway, K.A., Brotman, S.M., Rosen, J.D., Currin, K.W., Alkhawaja, A.A., Etheridge, A.S., Wright, F., Gallins, P., Jima, D., Zhou, Y., et al. (2024). Liver eQTL meta-analysis illuminates potential molecular mechanisms of cardiometabolic traits. The American Journal of Human Genetics. 10.1016/j.ajhg.2024.07.017.

24. MacParland, S.A., Liu, J.C., Ma, X.-Z., Innes, B.T., Bartczak, A.M., Gage, B.K., Manuel, J., Khuu, N., Echeverri, J., Linares, I., et al. (2018). Single cell RNA sequencing of human liver reveals distinct intrahepatic macrophage populations. Nat Commun 9, 4383. 10.1038/s41467-018-06318-7.

25. Aizarani, N., Saviano, A., Sagar, null, Mailly, L., Durand, S., Herman, J.S., Pessaux, P., Baumert, T.F., and Grün, D. (2019). A human liver cell atlas reveals heterogeneity and epithelial progenitors. Nature 572, 199–204. 10.1038/s41586-019-1373-2.

26. Payen, V.L., Lavergne, A., Alevra Sarika, N., Colonval, M., Karim, L., Deckers, M., Najimi, M., Coppieters, W., Charloteaux, B., Sokal, E.M., et al. (2021). Single-cell RNA sequencing of human liver reveals hepatic stellate cell heterogeneity. JHEP Rep 3, 100278. 10.1016/j.jhepr.2021.100278.

27. Andrews, T.S., Atif, J., Liu, J.C., Perciani, C.T., Ma, X.-Z., Thoeni, C., Slyper, M., Eraslan, G., Segerstolpe, A., Manuel, J., et al. (2022). Single-Cell, Single-Nucleus, and Spatial RNA Sequencing of the Human Liver Identifies Cholangiocyte and Mesenchymal Heterogeneity. Hepatol Commun 6, 821–840. 10.1002/hep4.1854.

28. Hong, S.E., Mun, S.J., Lee, Y.J., Yoo, T., Suh, K.-S., Kang, K.W., Son, M.J., Kim, W., and Choi, M. (2025). Single-cell eQTL analysis identifies genetic variation underlying metabolic dysfunction-associated steatohepatitis. Nat Genet, 1–11. 10.1038/s41588-025-02237-8.

29. Kim, H.Y., Rosenthal, S.B., Liu, X., Miciano, C., Hou, X., Miller, M., Buchanan, J., Poirion, O.B., Chilin-Fuentes, D., Han, C., et al. (2024). Multi-Modal Analysis of human Hepatic Stellate Cells identifies novel therapeutic targets for Metabolic Dysfunction-Associated Steatotic Liver Disease. J Hepatol, S0168-8278(24)02667-9. 10.1016/j.jhep.2024.10.044.

30. Taliun, D., Harris, D.N., Kessler, M.D., Carlson, J., Szpiech, Z.A., Torres, R., Taliun, S.A.G., Corvelo, A., Gogarten, S.M., Kang, H.M., et al. (2021). Sequencing of 53,831 diverse genomes from the NHLBI TOPMed Program. Nature 590, 290–299. 10.1038/s41586-021-03205-y.

31. Danecek, P., Bonfield, J.K., Liddle, J., Marshall, J., Ohan, V., Pollard, M.O., Whitwham, A., Keane, T., McCarthy, S.A., Davies, R.M., et al. (2021). Twelve years of SAMtools and BCFtools. Gigascience 10, giab008. 10.1093/gigascience/giab008.

32. Frankish, A., Diekhans, M., Ferreira, A.-M., Johnson, R., Jungreis, I., Loveland, J., Mudge, J.M., Sisu, C., Wright, J., Armstrong, J., et al. (2019). GENCODE reference annotation for the human and mouse genomes. Nucleic Acids Res. 47, D766–D773. 10.1093/nar/gky955.

33. Quinlan, A.R. (2014). BEDTools: The Swiss-Army Tool for Genome Feature Analysis. Curr Protoc Bioinformatics 47, 11.12.1–34. 10.1002/0471250953.bi1112s47.

34. Kang, H.M., Subramaniam, M., Targ, S., Nguyen, M., Maliskova, L., McCarthy, E., Wan, E., Wong, S., Byrnes, L., Lanata, C.M., et al. (2018). Multiplexed droplet single-cell RNA-sequencing using natural genetic variation. Nat Biotechnol 36, 89–94. 10.1038/nbt.4042.

35. Hao, Y., Stuart, T., Kowalski, M., Choudhary, S., Hoffman, P., Hartman, A., Srivastava, A., Molla, G., Madad, S., Fernandez-Granda, C., et al. (2024). Dictionary learning for integrative, multimodal, and massively scalable single-cell analysis. Nat Biotechnol 42, 293–304. 10.1038/s41587-023-01767-y.

36. Stuart, T., Srivastava, A., Madad, S., Lareau, C.A., and Satija, R. (2021). Single-cell chromatin state analysis with Signac. Nat Methods 18, 1333–1341. 10.1038/s41592-021-01282-5.

37. Yang, S., Corbett, S.E., Koga, Y., Wang, Z., Johnson, W.E., Yajima, M., and Campbell, J.D. (2020). Decontamination of ambient RNA in single-cell RNA-seq with DecontX. Genome Biol 21, 57. 10.1186/s13059-020-1950-6.

38. Korsunsky, I., Millard, N., Fan, J., Slowikowski, K., Zhang, F., Wei, K., Baglaenko, Y., Brenner, M., Loh, P.-R., and Raychaudhuri, S. (2019). Fast, sensitive and accurate integration of single-cell data with Harmony. Nat Methods 16, 1289–1296. 10.1038/s41592-019-0619-0.

39. Zhang, Y., Liu, T., Meyer, C.A., Eeckhoute, J., Johnson, D.S., Bernstein, B.E., Nusbaum, C., Myers, R.M., Brown, M., Li, W., et al. (2008). Model-based Analysis of ChIP-Seq (MACS). Genome Biology 9, R137. 10.1186/gb-2008-9-9-r137.

40. Amemiya, H.M., Kundaje, A., and Boyle, A.P. (2019). The ENCODE Blacklist: Identification of Problematic Regions of the Genome. Sci Rep 9, 9354. 10.1038/s41598-019-45839-z.

41. Orchard, P., Kyono, Y., Hensley, J., Kitzman, J.O., and Parker, S.C.J. (2020). Quantification, Dynamic Visualization, and Validation of Bias in ATAC-Seq Data with ataqv. Cell Syst 10, 298–306.e4. 10.1016/j.cels.2020.02.009.

42. Lee, S., Cook, D., and Lawrence, M. (2019). plyranges: a grammar of genomic data transformation. Genome Biol 20, 4. 10.1186/s13059-018-1597-8.

43. R Core Team (2015). R: A Language and Environment for Statistical Computing (R Foundation for Statistical Computing).

44. Liao, Y., Smyth, G.K., and Shi, W. (2014). featureCounts: an efficient general purpose program for assigning sequence reads to genomic features. Bioinformatics 30, 923– 930. 10.1093/bioinformatics/btt656.

45. Risso, D., Schwartz, K., Sherlock, G., and Dudoit, S. (2011). GC-Content Normalization for RNA-Seq Data. BMC Bioinformatics 12, 480. 10.1186/1471-2105-12-480.

46. Love, M.I., Huber, W., and Anders, S. (2014). Moderated estimation of fold change and dispersion for RNA-seq data with DESeq2. Genome Biology 15, 550. 10.1186/s13059-014-0550-8.

47. Benjamini, Y., and Hochberg, Y. (1995). Controlling the false discovery rate: a practical and powerful approach to multiple testing. Journal of the royal statistical society. Series B (Methodological) 57, 289–300.

48. McLean, C.Y., Bristor, D., Hiller, M., Clarke, S.L., Schaar, B.T., Lowe, C.B., Wenger, A.M., and Bejerano, G. (2010). GREAT improves functional interpretation of cis-regulatory regions. Nat. Biotechnol. 28, 495–501. 10.1038/nbt.1630.

49. The Gene Ontology Consortium (2019). The Gene Ontology Resource: 20 years and still GOing strong. Nucleic Acids Res. 47, D330–D338. 10.1093/nar/gky1055.

50. Heinz, S., Benner, C., Spann, N., Bertolino, E., Lin, Y.C., Laslo, P., Cheng, J.X., Murre, C., Singh, H., and Glass, C.K. (2010). Simple combinations of lineage-determining transcription factors prime cis-regulatory elements required for macrophage and B cell identities. Mol. Cell 38, 576–589. 10.1016/j.molcel.2010.05.004.

51. Bulik-Sullivan, B.K., Loh, P.-R., Finucane, H.K., Ripke, S., Yang, J., Schizophrenia Working Group of the Psychiatric Genomics Consortium, Patterson, N., Daly, M.J., Price, A.L., and Neale, B.M. (2015). LD Score regression distinguishes confounding from polygenicity in genome-wide association studies. Nat Genet 47, 291–295. 10.1038/ng.3211.

52. Hinrichs, A.S., Karolchik, D., Baertsch, R., Barber, G.P., Bejerano, G., Clawson, H., Diekhans, M., Furey, T.S., Harte, R.A., Hsu, F., et al. (2006). The UCSC Genome Browser Database: update 2006. Nucleic Acids Res. 34, D590–598. 10.1093/nar/gkj144.

53. Geijn, B. van de, McVicker, G., Gilad, Y., and Pritchard, J.K. (2015). WASP: allele-specific software for robust molecular quantitative trait locus discovery. Nat. Methods 12, 1061–1063. 10.1038/nmeth.3582.

54. Li, H. (2013). Aligning sequence reads, clone sequences and assembly contigs with BWA-MEM.

55. Ongen, H., Buil, A., Brown, A.A., Dermitzakis, E.T., and Delaneau, O. (2015). Fast and efficient QTL mapper for thousands of molecular phenotypes. Bioinformatics 32, 1479–1485.

56. Huang, L., Rosen, J.D., Sun, Q., Chen, J., Wheeler, M.M., Zhou, Y., Min, Y.-I., Kooperberg, C., Conomos, M.P., Stilp, A.M., et al. (2022). TOP-LD: A tool to explore linkage disequilibrium with TOPMed whole-genome sequence data. Am J Hum Genet 109, 1175–1181. 10.1016/j.ajhg.2022.04.006.

57. Yengo, L., Sidorenko, J., Kemper, K.E., Zheng, Z., Wood, A.R., Weedon, M.N., Frayling, T.M., Hirschhorn, J., Yang, J., Visscher, P.M., et al. (2018). Meta-analysis of genome-wide association studies for height and body mass index in ∼700000 individuals of European ancestry. Hum. Mol. Genet. 27, 3641–3649. 10.1093/hmg/ddy271.

58. Revez, J.A., Lin, T., Qiao, Z., Xue, A., Holtz, Y., Zhu, Z., Zeng, J., Wang, H., Sidorenko, J., Kemper, K.E., et al. (2020). Genome-wide association study identifies 143 loci associated with 25 hydroxyvitamin D concentration. Nat Commun 11, 1647. 10.1038/s41467-020-15421-7.

59. Said, S., Pazoki, R., Karhunen, V., Võsa, U., Ligthart, S., Bodinier, B., Koskeridis, F., Welsh, P., Alizadeh, B.Z., Chasman, D.I., et al. (2022). Genetic analysis of over half a million people characterises C-reactive protein loci. Nat Commun 13, 2198. 10.1038/s41467-022-29650-5.

60. Cerezo, M., Sollis, E., Ji, Y., Lewis, E., Abid, A., Bircan, K.O., Hall, P., Hayhurst, J., John, S., Mosaku, A., et al. (2025). The NHGRI-EBI GWAS Catalog: standards for reusability, sustainability and diversity. Nucleic Acids Research 53, D998–D1005. 10.1093/nar/gkae1070.

61. Ramachandran, P., Dobie, R., Wilson-Kanamori, J.R., Dora, E.F., Henderson, B.E.P., Luu, N.T., Portman, J.R., Matchett, K.P., Brice, M., Marwick, J.A., et al. (2019). Resolving the fibrotic niche of human liver cirrhosis at single-cell level. Nature 575, 512–518. 10.1038/s41586-019-1631-3.

62. Zhang, M., Yang, H., Wan, L., Wang, Z., Wang, H., Ge, C., Liu, Y., Hao, Y., Zhang, D., Shi, G., et al. (2020). Single-cell transcriptomic architecture and intercellular crosstalk of human intrahepatic cholangiocarcinoma. Journal of Hepatology 73, 1118–1130. 10.1016/j.jhep.2020.05.039.

63. Lim, Y.J., Koo, J.E., Hong, E.-H., Park, Z.-Y., Lim, K.-M., Bae, O.-N., and Lee, J.Y. (2015). A Src-family-tyrosine kinase, Lyn, is required for efficient IFN-β expression in pattern recognition receptor, RIG-I, signal pathway by interacting with IPS-1. Cytokine 72, 63–70. 10.1016/j.cyto.2014.12.008.

64. Musunuru, K., Strong, A., Frank-Kamenetsky, M., Lee, N.E., Ahfeldt, T., Sachs, K.V., Li, X., Li, H., Kuperwasser, N., Ruda, V.M., et al. (2010). From noncoding variant to phenotype via SORT1 at the 1p13 cholesterol locus. Nature 466, 714–719. 10.1038/nature09266.

65. Poisson, J., Lemoinne, S., Boulanger, C., Durand, F., Moreau, R., Valla, D., and Rautou, P.-E. (2017). Liver sinusoidal endothelial cells: Physiology and role in liver diseases. J Hepatol 66, 212–227. 10.1016/j.jhep.2016.07.009.

66. Marrone, G., Shah, V.H., and Gracia-Sancho, J. (2016). Sinusoidal communication in liver fibrosis and regeneration. J Hepatol 65, 608–617. 10.1016/j.jhep.2016.04.018.

67. Dekky, B., Azar, F., Bonnier, D., Monseur, C., Kalebić, C., Arpigny, E., Colige, A., Legagneux, V., and Théret, N. (2023). ADAMTS12 is a stromal modulator in chronic liver disease. FASEB J 37, e23237. 10.1096/fj.202200692RRRR.

68. Degner, J.F., Pai, A.A., Pique-Regi, R., Veyrieras, J.-B., Gaffney, D.J., Pickrell, J.K., De Leon, S., Michelini, K., Lewellen, N., Crawford, G.E., et al. (2012). DNase I sensitivity QTLs are a major determinant of human expression variation. Nature 482, 390–394. 10.1038/nature10808.

69. Patel, R., Nair, S., Choudhry, H., Jaffry, M., and Dastjerdi, M. (2024). Ocular manifestations of liver disease: an important diagnostic aid. Int Ophthalmol 44, 177. 10.1007/s10792-024-03103-y.

70. Chen, M.-H., Raffield, L.M., Mousas, A., Sakaue, S., Huffman, J.E., Moscati, A., Trivedi, B., Jiang, T., Akbari, P., Vuckovic, D., et al. (2020). Trans-ethnic and Ancestry-Specific Blood-Cell Genetics in 746,667 Individuals from 5 Global Populations. Cell 182, 1198–1213.e14. 10.1016/j.cell.2020.06.045.

71. Vuckovic, D., Bao, E.L., Akbari, P., Lareau, C.A., Mousas, A., Jiang, T., Chen, M.-H., Raffield, L.M., Tardaguila, M., Huffman, J.E., et al. (2020). The Polygenic and Monogenic Basis of Blood Traits and Diseases. Cell 182, 1214–1231.e11. 10.1016/j.cell.2020.08.008.

72. Shen, Z., Shen, B., Dai, W., Zhou, C., Luo, X., Guo, Y., Wang, J., Xu, X., Sun, Z., Cai, X., et al. (2023). Expansion of macrophage and liver sinusoidal endothelial cell subpopulations during non-alcoholic steatohepatitis progression. iScience 26, 106572. 10.1016/j.isci.2023.106572.

73. Suzuki, J., Imanishi, E., and Nagata, S. (2014). Exposure of Phosphatidylserine by Xk-related Protein Family Members during Apoptosis. Journal of Biological Chemistry 289, 30257–30267. 10.1074/jbc.M114.583419.

74. Bárcena, C., Stefanovic, M., Tutusaus, A., Joannas, L., Menéndez, A., García-Ruiz, C., Sancho-Bru, P., Marí, M., Caballeria, J., Rothlin, C.V., et al. (2015). Gas6/Axl pathway is activated in chronic liver disease and its targeting reduces fibrosis via hepatic stellate cell inactivation. J Hepatol 63, 670–678. 10.1016/j.jhep.2015.04.013.

75. Staufer, K., Huber, H., Zessner-Spitzenberg, J., Stauber, R., Finkenstedt, A., Bantel, H., Weiss, T.S., Huber, M., Starlinger, P., Gruenberger, T., et al. (2023). Gas6 in chronic liver disease-a novel blood-based biomarker for liver fibrosis. Cell Death Discov 9, 282. 10.1038/s41420-023-01551-6.

